# Insight parameter drug design for human β-tryptase inhibition integrated molecular docking, QSAR, molecular dynamics simulation, and pharmacophore modelling studies of α-keto-[1,2,4]-oxadiazoles

**DOI:** 10.1101/2022.07.17.500327

**Authors:** Chai Xin Yu, Jian Wei Tan, Kamal Rullah, Syarul Imran, Chau Ling Tham

**Affiliations:** Department of Biomedical Science, Faculty of Medicine and Health Sciences, Universiti Putra Malaysia, 43400 UPM Serdang, Selangor, Malaysia; Drug Discovery and Synthetic Chemistry Research Group, Department of Pharmaceutical Chemistry, Kulliyyah of Pharmacy, International Islamic University Malaysia, Bandar Indera Mahkota, 25200 Kuantan, Pahang, Malaysia; Atta-ur-Rahman Institute for Natural Product Discovery (QuRIns),Universiti Teknologi MARA Cawangan Selangor Kampus Punca Alam, 42300 Puncak Alam, Selangor, Malaysia; Faculty of Applied Science, Universiti Teknologi MARA (UiTM), 40450 Shah Alam, Selangor, Malaysia

**Keywords:** β-tryptase inhibitor, 2D-QSAR, molecular docking, molecular dynamics simulations, pharmacophore modelling

## Abstract

Dengue hemorrhagic fever (DHF) is life-threatening severe dengue with a hallmark of vascular leakage. A mast cell protease, β-tryptase, has been found to promote vascular leakage in DHF patients, which could be a potential target for the treatment of DHF. This study aims to develop a theoretical background for the design and selection of β-tryptase inhibitors through these approaches: two-dimensional quantitative structure-activity relationships (2D-QSAR) study, molecular docking analysis, molecular dynamics (MD) simulation, and structure-based pharmacophore modelling (PM). A total of 34 a-keto-[1,2,3]-oxadiazoles scaffold-based compounds, obtained from a past study, were used to generate 2D-QSAR models by Genetic Function Approximation (GFA). The generated 2D-QSAR models were used to investigate the relationships between the molecular structure and the potency of β-tryptase inhibition. Molecular docking explores the binding affinities and binding interactions of the a-keto-[1,2,3]-oxadiazoles scaffold-based compounds with β-tryptase (PDB Code 4A6L) by the CDOCKER tool in Discovery Studio. In addition, MD simulation was performed using GROMACS on the docked complex of the reported most active compound, compound 11e, to study the binding mechanism of the compound towards β-tryptase. Finally, a structure-based pharmacophore model was generated from the same docked complex to identify the important features that contribute positively to the inhibitory activity of the compound towards β-tryptase. The best 2D-QSAR model has demonstrated statistically significant results through its r^2^, q^2^, and r^2^ (pred) values of 0.9077, 0.733, and 0.8104, respectively. The docking results of compound 11e showed lower CDOCKER energy than the 4A6L co-crystallised ligand, indicating good binding affinity. Furthermore, compound 11e has a similar binding pattern as the 4A6L co-crystallised ligand, which involves the binding of active residues such as Asp207, SER208, and GLY237. The MD simulation shows that the 4A6L-compound 11e complex has RMSD below 2Å throughout the 100ns of simulation, indicating that the docked complex is stable. Besides, MD simulation showed that the inhibitory potency of compound 11e is contributed by hydrogen bonding with 4A6L active site residues, which are ASP207, SER208, and GLY237. The best pharmacophore model identified features that contribute to the inhibitory potency of a compound, which included hydrogen bond acceptor, ionic interaction, hydrophobic interaction, and aromatic ring. This study has fundamentally supplied valuable insight and knowledge for developing novel chemical compounds with improved inhibitory ability against human β-tryptase.

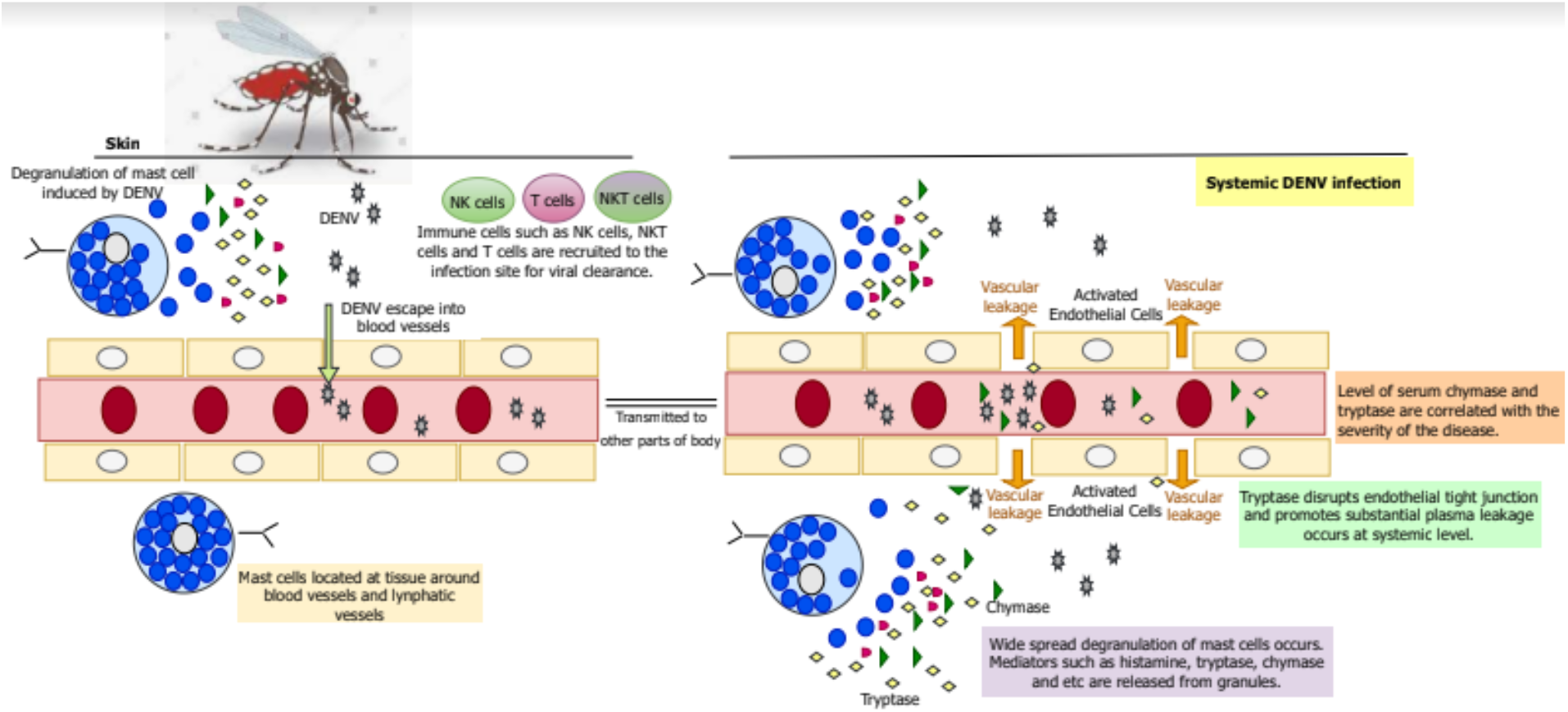

DENV induces mast cell degranulation to release mediators, such as β-tryptase. β-Tryptase has been reported to promote plasma leakage, which is the hallmark of dengue hemorrhagic fever (DHF). Thus, β-tryptase could be a promising target for the treatment of DHF.

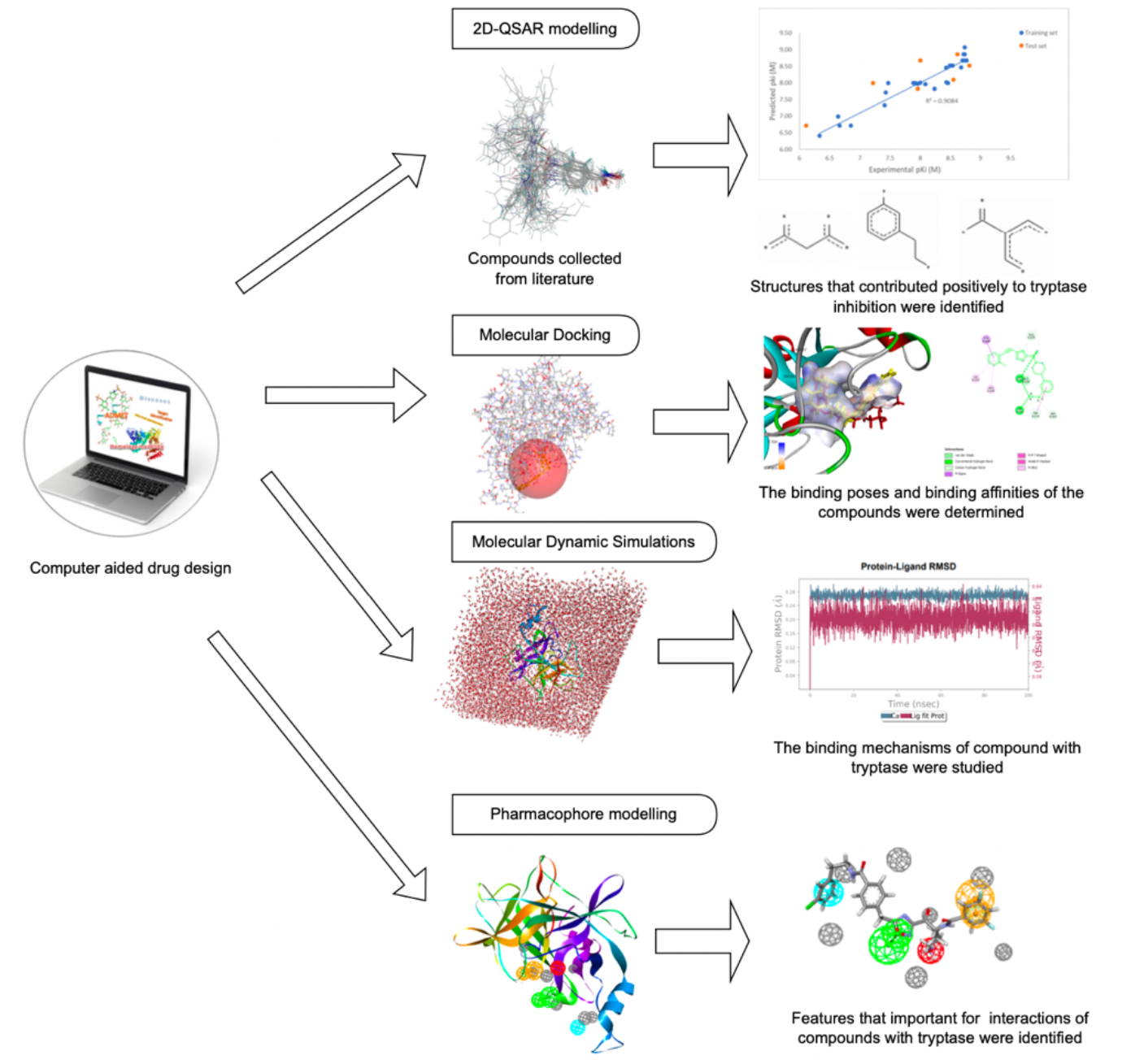

In this study, computer-aided drug design was used to examine the structure-activity relationship and binding mechanisms of β-tryptase inhibitors reported in a past study. Finally, theoretical background for the design of β-tryptase inhibitors was built.

## 1. Introduction

Dengue is a mosquito-transmitted viral infection that commonly occurs globally, and around half of the global population lives in a region where dengue is regularly found[1][2]. Infections occur each year in the estimation of 390 million cases. Among these cases, 96 million are with clinical manifestation [1]–[3]. A total of 129 countries are at risk of infection, with the highest burden in Asia [2]. Dengue virus (DENV) is a flavivirus transmitted by Aedes mosquitoes, mainly by *Aedes aegypti* and to a lesser extent by *Aedes Albopictus* [2]. Patients with primary or initial secondary dengue infection will have the mild-acute undifferentiated febrile illness to classical Dengue Fever (DF). Some patients develop dengue hemorrhagic fever (DHF) which is characterised by vascular leakage [4]. This vascular leakage can result in dengue shock syndrome (DSS) with an imperceptible pulse and is usually fatal [4].

The primary prevention method of dengue is vector control, while medications and effective vaccines for dengue are still under development. At the time of writing, Dengvaxia® (Sanofi Pasteur) is the only dengue vaccine that has been approved for use in people over nine years old in around 20 countries worldwide since it was first licensed in Mexico in 2015 [5], [6]. However, the efficacy of Dengvaxia® vaccination is varied across different serotypes, ages, and status of dengue sera before vaccination [5], [6]. Besides, the WHO Strategic Advisory Panel has recommended the use of Dengvaxia® for dengue-seropositive patients only because it has been found to increase the risk of severe dengue in seronegative individuals [5], [6]. To date, dengue treatment has relied on the management of clinical signs and symptoms, including the prescription of non-steroidal anti-inflammatory agents (NSAIDs), clinical parameters tracking, and fluid management to stabilise hemodynamic status [7]. Currently, there is still no specific treatment for dengue [2], [7], [8]. Although some drugs such as chloroquine, balapiravir, and lovastatin are being tested in human clinical trials to treat dengue at different stages of development, these drugs have not been approved yet for use against dengue [9], [10].

DENV has been found to trigger protective immunological responses and mast cell (MC) degranulation in mouse models [11], [12]. When mosquitoes inject DENV into the host body, the degranulation of MCs and the release of mediators such as histamine, chymase, tryptase and etc will be triggered. These mediators increase the permeability of blood vessels and recruit immune cells to the infection site for viral clearance [12], [13]. However, when local protection mechanisms fall short, DENVs enter the bloodstream and are transported to other parts of the body, which cause systemic infection and widespread activation of MCs degranulation [12], [13]. [11]A few studies found an association between the clinical sign of DHF and MCs activation [11], [14], [15]. MCs specific products are found high in the blood and urine of human DENV patients [11]. In another study, [12] also reported that in response to DENV, DENV-induced vascular leakage was not observed in MCs-deficient mice as well as in wild-type mice treated with clinically approved MCs stabilising compound. These findings have suggested MCs play an important role in DENV-induced vascular leakage. Tryptase has been reported to promote substantial vascular leakage that could cause vascular pathology by reducing the expression of endothelial cell adhesion molecules and breaking endothelial tight junctions [11]. The level of tryptase is found to be associated with the grade of DHF severity in independent human dengue cohorts [11]. Thus, tryptase could be a potential target for the treatment of DENV-induced vascular leakage [11].

Human tryptase is a trypsin-like protein and it is the most abundant protein in MCs [16]–[18]. There are different types of tryptase in the granules of human MCs and β-tryptase is the main form of tryptase that is stored as an active enzyme in MCs [16], [19]. β-Tryptase is a tetramer unit. The active site of each monomer faces one another and forms a flat rectangular frame with a narrow opening, around 50Å x 30Å at the center [18], [20], [21]. This limitation has caused difficulty in the development of β-tryptase inhibitors as only small, conformationally flexible molecules or proteins are able to access the active site cavity. Numerous compounds have been studied to inhibit tryptase. However, the development of tryptase inhibitors has faced problems such as safety issues, poor bioavailability, low potency, and selectivity issues [17], [22], [23]. Nafamostat Mesylate was found to be a very potent β-tryptase inhibitor, however, it has low selectivity over other trypsin-like protease at a higher concentration and it inhibits β-tryptase by suicide inhibition [24], [25]. Most of the β-tryptase inhibitors have limited success in clinical trials and none of the β-tryptase inhibitors has successfully passed clinical trials to date [23], [26]. Therefore, research to identify drugs potentially inhibit tryptase, particularly for the treatment of DENV-induced vascular leakage caused by tryptase, remains paramount.

Computer-aided drug design (CADD) is a powerful tool for improving the efficiency of the drug discovery and development process [27]. The example of CADD methods in drug design included quantitative structure-activity relationship (QSAR), molecular docking, molecular dynamics simulation (MD), and pharmacophore modelling (PM). QSAR is a method to quantify the relationship between structure and activity as well as to provide information on how structure affects activity. Approaches in QSAR included 2D-QSAR and 3D-QSAR [28]. On the one hand, molecular docking predicts binding poses and binding affinity of a ligand to a biological molecule, while MD simulation prostrates the binding mechanisms of the ligand to the biological molecule at a microscopic level [29], [30]. On the other hand, PM in the drug design-construct hypothesis helps to understand features that support interactions between the ligand and the biological molecule [31].

A total of 34 α-keto-[1,2,4]-oxadiazoles scaffold-based derivatives with inhibitory activity against human β-tryptase have been reported by [32]. The compounds reported have high potency and selectivity towards β-tryptase. However, the authors have not examined the compounds’ structure-activity relationship and binding-based inhibition mechanisms. Hence, our present study aims to identify the QSAR and the binding affinities of this series of α-keto-[1,2,4]-oxadiazoles scaffold-based compounds with human β-tryptase. The binding mode of the α-keto-[1,2,4]-oxadiazoles scaffold-based compounds with β-tryptase was also investigated for the construction of theoretical background. In this study, 2D-QSAR was used to determine the relationship between α-Keto-[1,2,4]-oxadiazoles scaffold-based compounds and their activities toward β-tryptase inhibition. Besides, the binding affinities and binding poses of α-keto-[1,2,4]-oxadiazoles scaffold-based compounds with β-tryptase were studied using molecular docking. The interactions between the most active compound with β-tryptase were investigated by MD. Finally, a model was constructed using the structure-based pharmacophore modelling based on the protein-compound complex obtained from the last frame of MD. Our findings from this study may help in the design and selection of β-tryptase inhibitors for the treatment of DENV-induced vascular leakage in the future.

## 2. Materials and Methods

### 2.1 Selection of data set

A total of thirty-four α-keto-[1,2,4]-oxadiazoles scaffold-based derivatives with inhibitory activities against human β-tryptase were collected from [32] for this study [33] (Table S1). The 2D structure of all the compounds was constructed using ChemDraw Professional 15.0 (Perkin Elmer Inc., Waltham, MA, USA), and all the compounds were converted to the 3D conformers structure by saving them as Discovery Studio Files in Discovery Studio^®^ 3.1 (Accelrys Inc. San Diego, CA, USA) (DS). The compounds were then prepared by standardising charges for common groups, adding hydrogens, enumerating ionisation states, ionising functional groups, removing duplicates, and generating 3D conformations [33]. Next, the structures were optimised by energy minimisation with the CHARMm force field in DS, and these compounds were subjected to 2D-QSAR and molecular docking analysis. Finally, the best-selected compound was further investigated in MD simulation and pharmacophore modelling. The overview of the methodology is shown in Figure 1. The 2D-QSAR study, molecular docking, and pharmacophore modelling were performed using DS, while MD simulation was done in Desmond.

**Figure 1:**
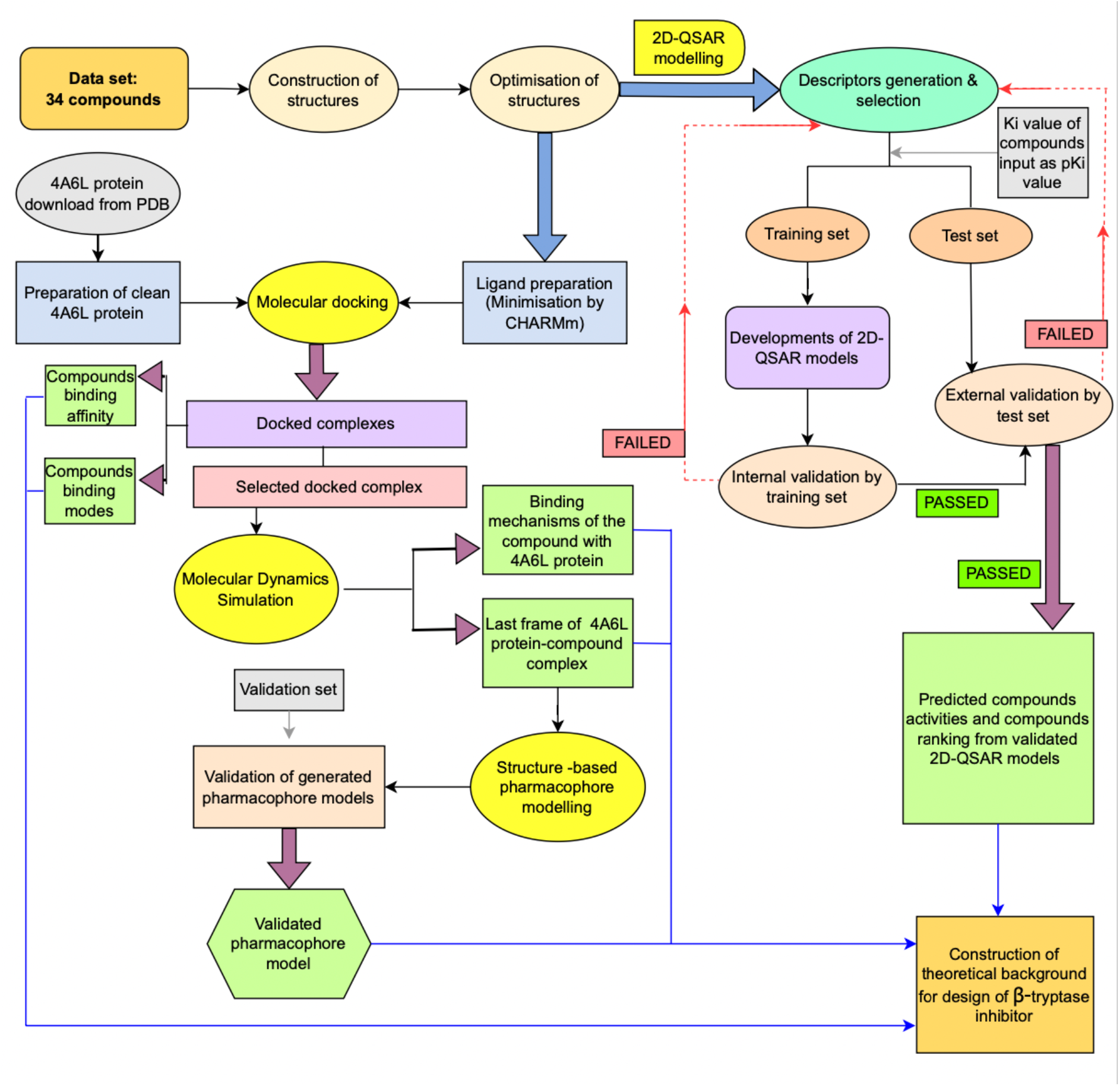
Flow chart for integrated molecular docking, 2D-QSAR modelling, MD simulation, and pharmacophore modelling. Results obtained from molecular docking, 2D-QSAR modelling, MD simulation, and pharmacophore modelling were used to construct theoretical background for the design of β-tryptase inhibitor.

### 2.2 Molecular docking

Docking studies were carried out using CDOCKER, the docking algorithms’ CHARMM program, in DS. Crystal structure of β-tryptase receptor in complex with ligand (1-{3-[1-({5-[(2-fluorophenyl)ethynyl]furan-2-yl} carbonyl)Piperidin-4-yl]phenyl}methanamine) was obtained from Protein Data Bank (PDB) with ID of 4A6L. The 4A6L is a tetramer with four identical monomers with the same amino acid sequence. The chain A monomer was selected for the docking study. In preparation for clean chain A, water molecules and ligand were removed using USFC Chimera version 1.14 (University of California, SF, USA) [34]–[36]. The structure was prepared by inserting missing atoms, modelling missing loop regions, optimising sidechain conformations, deleting alternate conformations, standardising atom names, and protonating titratable residues [33], [37]. Then, a binding-site sphere was defined at the structure of where the native ligand was bound. Validation of the docking method was performed by re-docking the native ligand into the defined binding site to reproduce the binding pose of the native ligand in 4A6L chain A observed in the crystal structure [38]. Root mean square deviation (RMSD) value was used to assess the difference between the redocked and the original pose among the top-ranking conformation clusters from the dock. The method that produced the docked result with a low RMSD value (below 2.0Å) was applied for the docking of α-keto-[1,2,4]-oxadiazoles scaffold-based compounds [38].

In the docking of α-keto-[1,2,4]-oxadiazoles scaffold-based compounds, the prepared protein 4A6L chain A was defined as the total receptor, and the active site was selected based on the ligand-binding region of the bound native ligand. After that, the native ligand was removed. The prepared dataset of the thirty-four compounds was used for the docking [39]. During docking, the 4A6L chain A structure was kept rigid while the ligands acted as flexible. Ten ligand-binding poses were generated and ranked according to their CDOCKER energy from the dock, and the predicted binding modes were analysed[39].

### 2.3 2D-QSAR modelling

#### 2.3.1 Preparation of compounds

The thirty-four compounds prepared at 2.1 with inhibitory activities predicted under the same biological assay conditions were applied to generate and validate 2D-QSAR models. The compounds and their inhibition constant (Ki) values were exported to DS. The Ki of these compounds were converted to a negative logarithmic (-logKi or pKi) scale, as shown in the supplementary document (Table S1) [32]. Then, the data set was divided into a training set and a test set. The full ligand set is sorted through a random index. The “Top N” ligands were then assigned to the training set based on the value of the Training Set Percentage parameter and inhibition constant of the compounds [40], [41]. The training set contained twenty-seven compounds, and the test set contained seven compounds. The training set was used to build the 2D-QSAR models, while the test set was used to access the performance of the generated models.

#### 2.3.2 Descriptors calculation and selection

The molecular properties of the thirty-four compounds were calculated, and descriptors included fingerprint and numeric properties that explained the biological activities of the compounds were identified based on the correlation with the pKi value [40]. Example of descriptors is AlogP, Extended Connectivity Fingerprint (ECFP_6), Molecular_Weight, Num_Aromatic Rings, Num_H_Acceptors, Num_H_Donors, Num_Rings, Num_Rotatable Bonds, Molecular_Fractional, Polar Surface Area and Topology Descriptors which included CHI, Kappa, etc. [41].

#### 2.3.3 Development of QSAR models and validation

The 2D-QSAR models were developed by Genetic Function Approximation (GFA) algorithms with the training set compounds using pKi values as the dependent variable and the calculated descriptors as independent variables. The quality of the models was examined by Friedman lack-of-fit (LOF), S.O.P. p-value, coefficient of determination (r^2^), adjusted r^2^ (adj-r^2^), r^2^(Pred), and cross-validated correlation coefficient (q^2^) [40]–[42].

### 2.4 Molecular dynamics simulation

MD simulation was performed on the selected docked complex by using Desmond. OPLS3e force field was applied to the docked complex and the system was solvated with a water molecule in a truncated octahedron box at a 10 Å distance. The system was run in periodic boundary conditions. First, the system was minimised with 2000 steps of steepest descent minimisation followed by 3000 steps of conjugate gradient minimisation. Then, the system was heated from temperature 0 to 300 K for 100 ns in the NVT ensemble. After that, the system was equilibrated in an NPT ensemble under the constant pressure of 1.0 bar for 100 ns. Finally, MD simulation was performed on the system under NPT ensemble conditions by using periodic boundary conditions and particle-mesh Ewald (PME) for long-range electrostatics [43].

### 2.5 Structure-based pharmacophore modelling

The last frame of protein 4A6L chain A-compound complex from MD was subjected to generate pharmacophore models using the Receptor-ligand Pharmacophore Generation protocol of DS. In this protocol, the pharmacophore models were generated from the ligand features that match the receptor-ligand interactions, including hydrogen bond acceptor, hydrogen bond donor, hydrophobic feature, ionisable feature, and aromatic ring [33]. Excluded volumes were added based on the locations of atoms on the protein. The top models were produced based on sensitivity and specificity, and the model with the highest selectivity, as predicted by Genetic Function Approximation (GFA) model, was selected. The selected pharmacophore model was edited by Edit and Cluster tool in DS to produce a final model with key pharmacophoric features [39], [44].

After that, the selected pharmacophore model was validated by a validation set that consisted of 81 active compounds (Ki/IC_50_ < 50nM), 64 inactive compounds (Ki/IC_50_ ≥ 1000 nM), and 7,860 compounds as decoys [38]. The active and inactive compounds were identified from past studies on β-tryptase inhibitors, and the compounds obtained from Chemdraw were optimised and minimised using DS. On the other hand, the decoy was prepared from small molecules downloaded from the ZINC database, followed by screening with Decoy Finder software [45]. The decoy molecules were also optimised and minimised using DS before being combined with the prepared active and inactive compounds.

The validation set was screened by the selected pharmacophore model using Ligand Pharmacophore Mapping Protocol with the best flexible conformation search method in DS [44]. The relevant ligand-pharmacophore mappings were exported and aligned to the pharmacophore. The accuracy, sensitivity, and specificity of the models were calculated as follows:

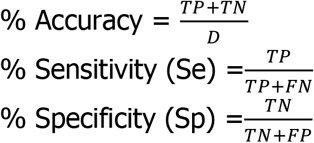

Whereby;

True positive (TP) is the number of compounds correctly identified as active
True negative (TN) is the number of compounds correctly identified as inactive
False-negative (FN) is the number of compounds incorrectly identified as inactive
False-positive (FP) is the number of compounds incorrectly identified as active
D is the total number of compounds in the validation set

The receiver operating characteristics (ROC) curve was generated, and the area under the curve (AUC) was determined [46]. Percentage yields of active (Y%), enrichment factor (EF), and Güner-Henry (GH) score, which envisioned the goodness of hit of the pharmacophore model screening, were calculated as follows [38]:

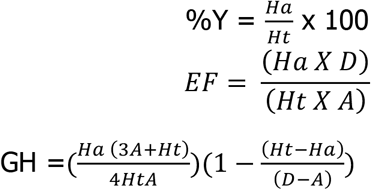

Whereby;

A is the total number of active compounds,
Ht is the total number of compounds screened by a pharmacophore model
Ha is the total number of active compounds obtained from screening.

## 3. Result and discussion

### 3.1 Molecular docking

> In the validation of the docking method, the pose of the native ligand before docking is similar to its docked pose with an RMSD value of 1.7Å, which is less than 2.0Å [37]. This showed that the CDOCKER method is applicable for molecular docking of the 4A6L chain A crystal structure in this study. The native ligand was used as a reference for the docking study. The interaction profile of the native ligand with 4A6L chain A is presented in Figure 2a. A total of thirty-four compounds were docked against 4A6L, which the docking result is presented in the Supplementary document (Table S2). A number of the compounds after docking have lower CDOCKER energy than the native ligand of −23.9672kJ/mol, including compound **11e** (N-(6-amino-1-(5-(4-((3-chlorophenethyl)carbomoyl)benzyl)-1,2,4-oxadiazol-3-yl)-1-oxohexan-2-yl)-3,5-difluorobenzamide) with CDOCKER energy of −40.9846KJ/mol and compound **1a** (benzyl (6-amino-1-(5-(4-methoxybenzyl)-1,2,4-oxadiazol-3-yl)-1-oxohexan-2-yl)carbamate) with CDOCKER energy of −36.3865kJ/mol, indicating that they have better binding affinities than the native ligand. Among all, compound **11e** is the most active compound, with a reported experimental activity of 1.5nM and high selectivity over trypsin [32]. The interaction profile of compound **11e** is shown in Figure 2b. On the other hand, compound **1a** has a reported experimental activity on β-tryptase of 770 nM, which is the most inactive among all compounds [32]. The interaction profile of compound **1a** is shown in Figure 2c.

**Figure 2:**
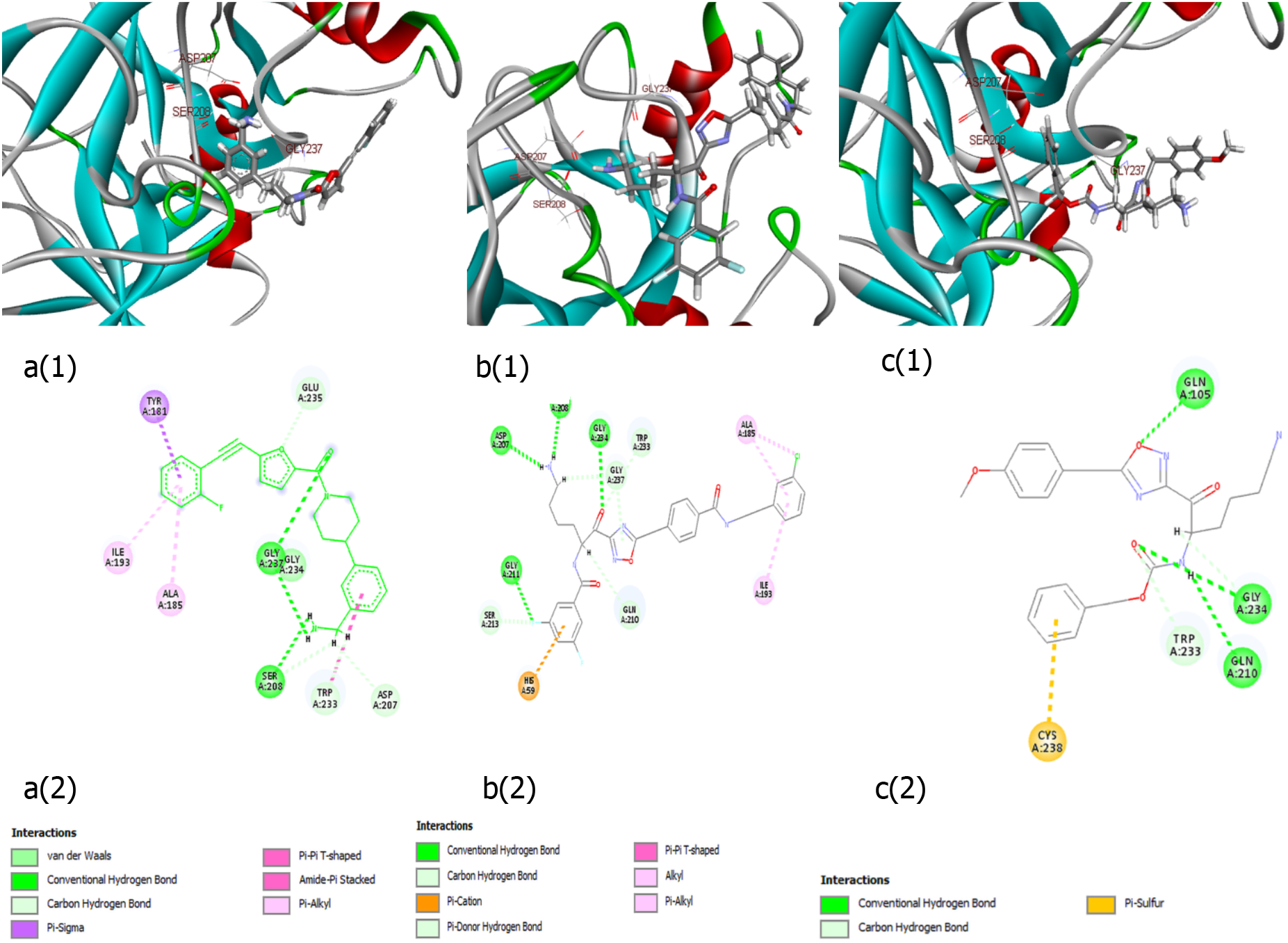
X-ray crystal structures and docked poses after molecular docking of (a)4A6L native ligand, (b)compound 11e, and (c)compound 1a

The binding pattern of compound **11e** was similar to the native ligand. Both native ligand and compound **11e** formed conventional hydrogen bonds with Asp207, Ser208, and Gly237, which are catalytic residues of S1 pocket in β-tryptase reported from past studies [21], [47]–[49]. The amino groups in compound **11e** and the native ligand bound to the protein residues Asp207 and Ser208 in a similar way. Besides, the carbonyl group of both the native ligand and compound 11e is also bound to protein residue Gly237 similarly. On the contrary, the binding pattern of compound **1a** is distinct from the native ligand, which differs in the types of bonding as well as the interacting atoms and protein residues. Overall, the molecular docking results from this study are consistent with [32].

### 3.2 2D-QSAR model

From the 2D-QSAR analysis on the α-keto-[1,2,4]-oxadiazole derivatives by using GFA, a total of ten 2D-QSAR equations were generated. Three 2D-QSAR equations were found to have robust statistical values and good correlations between physicochemical descriptors and biological activities (Table 1). The inhibitory activities of these compounds predicted by the three selected 2D-QSAR models are presented in the supplementary document (Table S3).

**Table 1:**
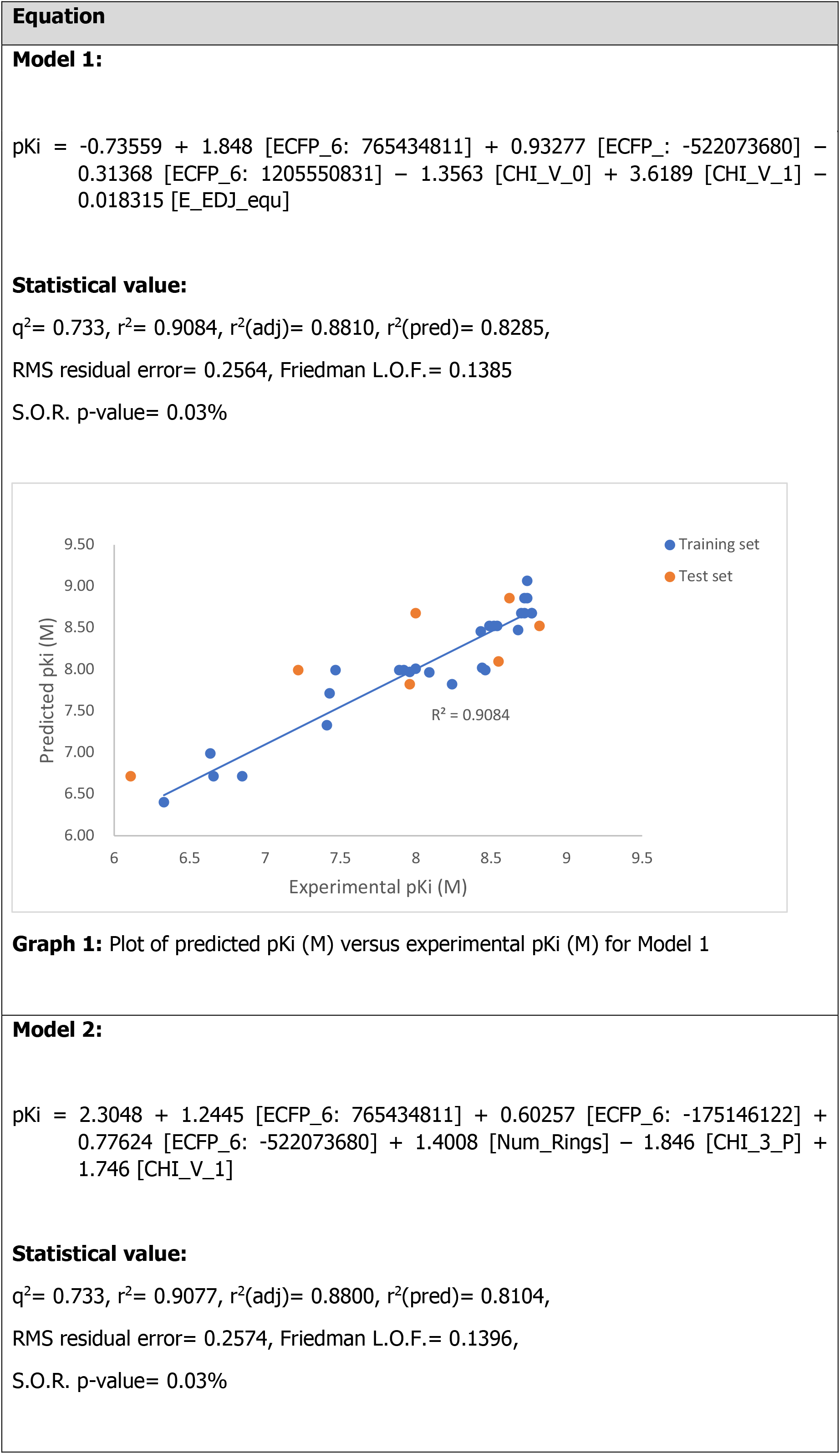
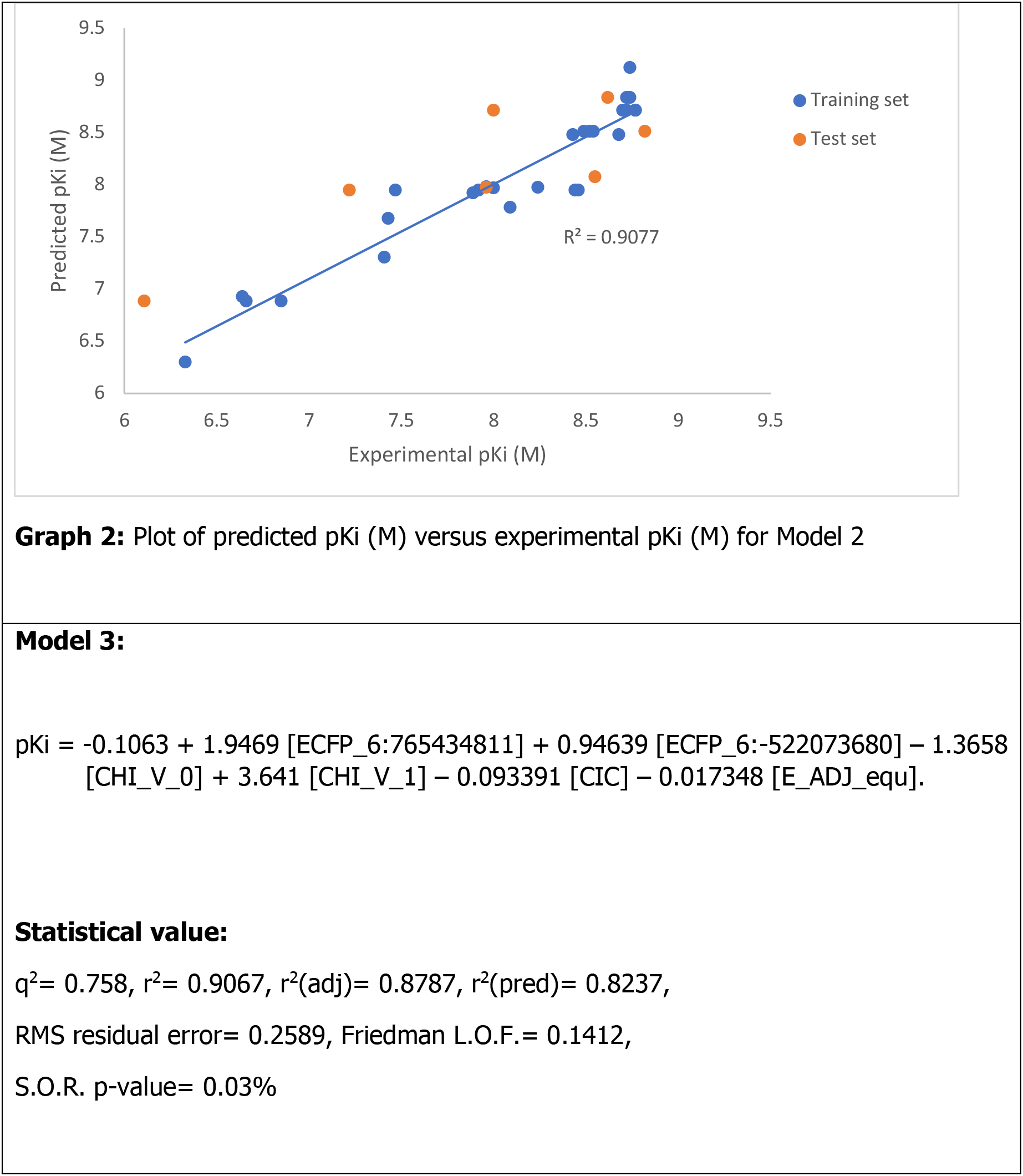
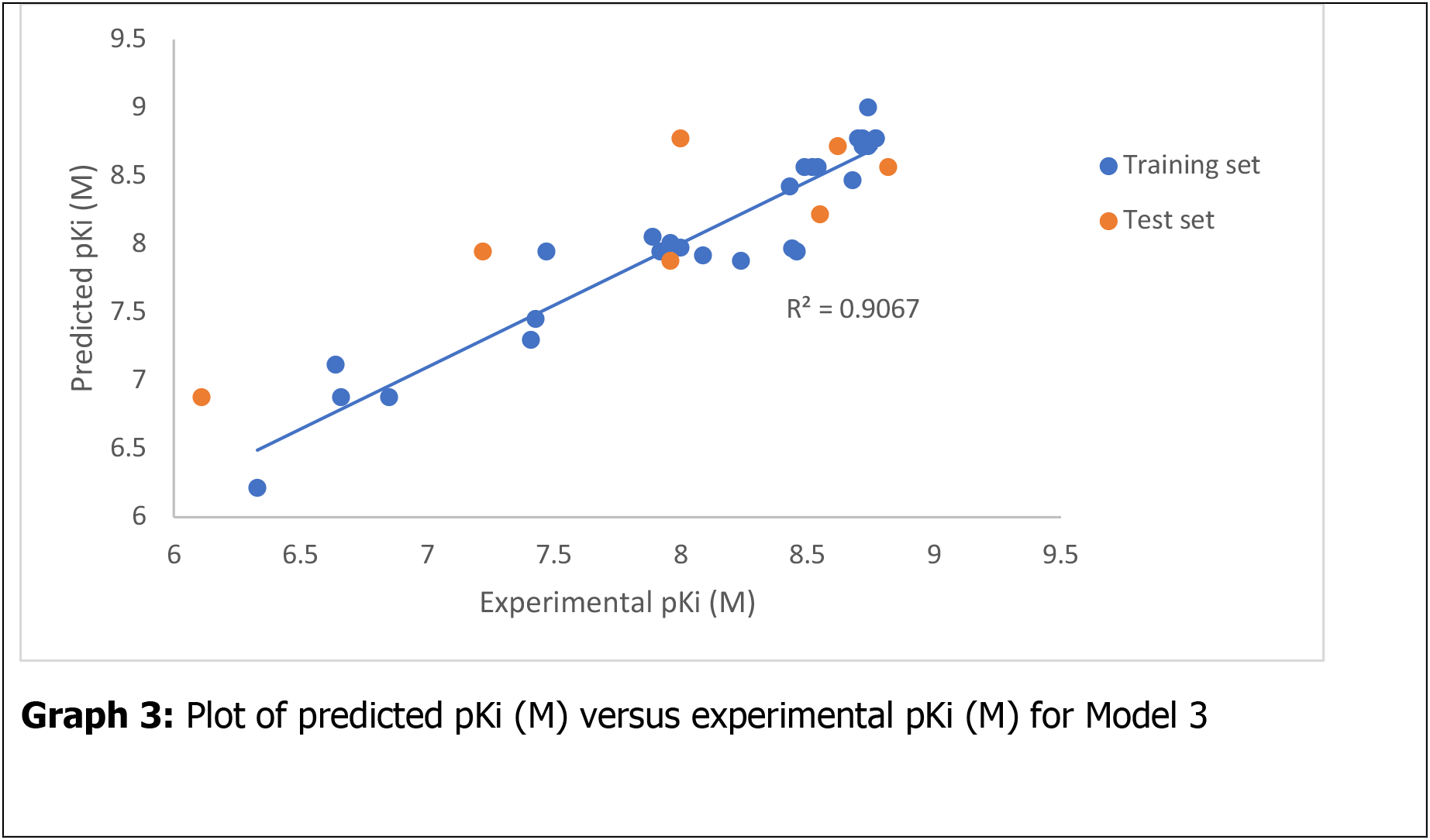
Results of the three best 2D-QSAR models

From Table 1, r^2^ is the squared correlation coefficient which indicates the agreement between the predicted and observed values of the training set [50], [51]. On the other hand, r^2^(adj) is the adjusted-square-correlation coefficient to determine the predictability of the model, while r^2^(pred) shows the complementary potential of the developed model [42]. The values of r^2^, r^2^(adj), and r^2^(pred) of the three selected models are all in the range of 0.81 to 0.91, which are higher than the minimum value of 0.8 for r^2^ and 0.6 for r^2^(adj) and r^2^(pred) [41], [51]. Besides, q^2^ reveals the predictability of the model [50]. The higher the q^2^ value, the stronger the fitting ability of the model [50]. The three developed models have q^2^ values between 0.733 to 0.758, which are more than the minimum value of 0.5. From the statistical analysis, the three developed models have shown strong relationships between selected structure features and the compounds’ biological activities.

Apart from that, all three developed models with RMS residual error, Friedman L.O.F., and S.O.P. p-value within the theoretical minimum value. RMS residual errors are the probability of obtaining the constant variance of values from a built model. A model with low RMS residual error has good repeatability, and the minimum value for RMS residue is between 0 to 1 [41], [51]. The RMS residual error of the three selected models is between 0.2564 to 0.2589. On the other hand, a model that resits overfitting has a Friedman L.O.F. value between 0 to 1. The Friedman L.O.F. value of the three developed models is between 0.1385 to 0.1412. A lower S.O.P. p-value indicates that the model is better, and the value should be within 0 to 5% [41]. The S.O.P. p-value of the three selected models is 0.03%. These show that the three developed models have good repeatability and good resistance to overfitting, and are statistically proven good models.

A total of ten molecular descriptors have been identified from the three selected models. The established correlation between descriptors and pKi values of the three models is explained in Table 2. In brief, the descriptors [ECFP_6: 765434811], [ECFP_6:-522073680], and [ECFP_6:-175146122] positively contributed to the increase of inhibitory activity of a compound towards β-tryptase. These descriptors could be found in the structures of some tryptase inhibitors reported in past studies (Table S3) [22], [24], [47], [49], [52]–[57].

**Table 2:**
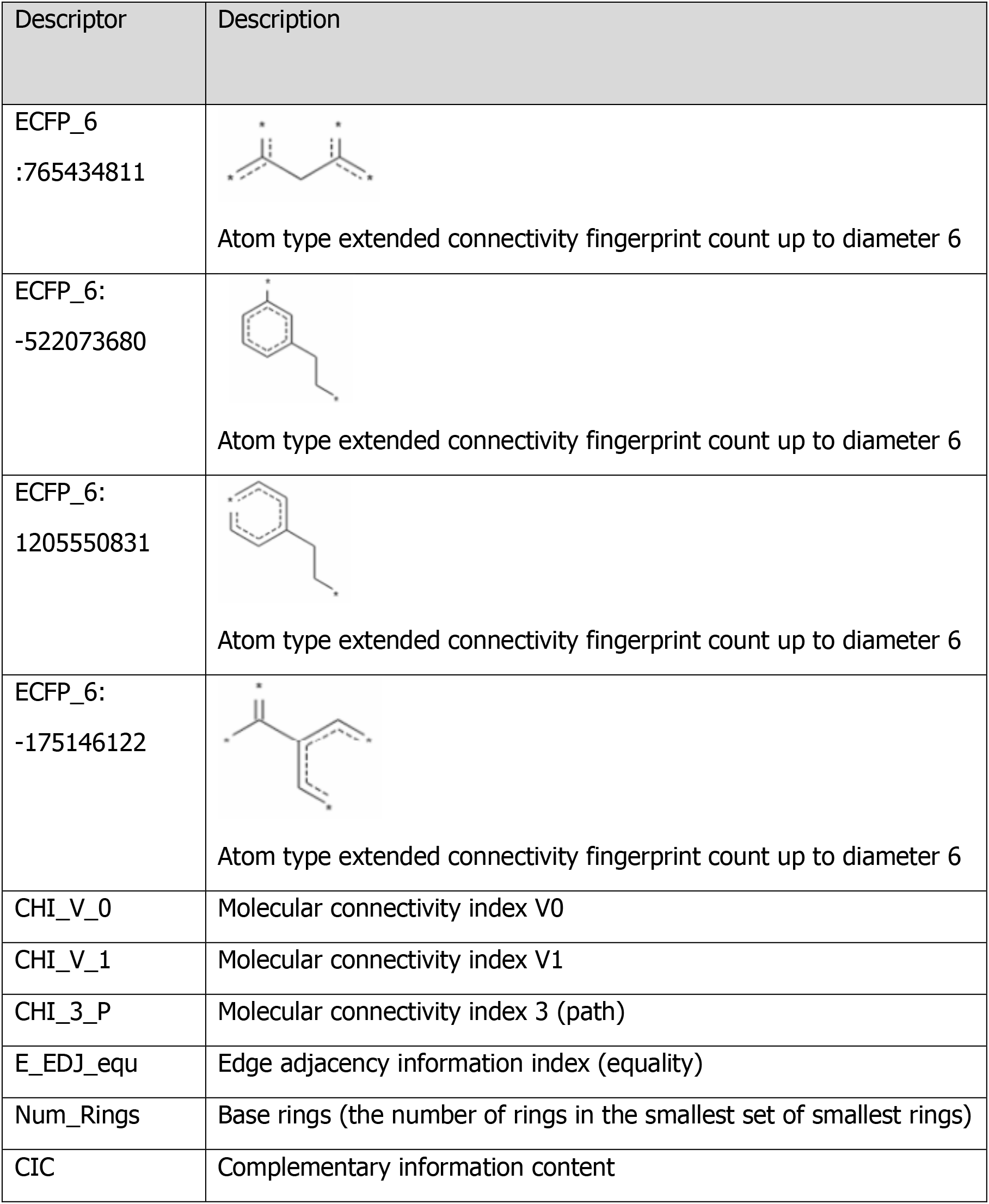
Descriptors identified from generated 2D-QSAR equations

| Descriptor | Description |
| --- | --- |
| ECFP_6<br>:765434811 | Atom type extended connectivity fingerprint count up to diameter 6 |
| ECFP_6:<br>-522073680 | Atom type extended connectivity fingerprint count up to diameter 6 |
| ECFP_6:<br>1205550831 | Atom type extended connectivity fingerprint count up to diameter 6 |
| ECFP_6:<br>-175146122 | Atom type extended connectivity fingerprint count up to diameter 6 |
| CHI_V_0 | Molecular connectivity index V0 |
| CHI_V_1 | Molecular connectivity index V1 |
| CHI_3_P | Molecular connectivity index 3 (path) |
| E_EDJ_equ | Edge adjacency information index (equality) |
| Num_Rings | Base rings (the number of rings in the smallest set of smallest rings) |
| CIC | Complementary information content |

Moreover, the descriptors [CHI_V_0], [CHI_V_1], and [CHI_3_P] are molecular connectivity indexes. [CHI_V_0] is a zero-order valence molecular connectivity index. The presence of long length and high branches of hydrocarbon chains will increase the value of this index while the value decreases with the presence of heteroatoms [58]. The zeroth-order valence molecular connectivity index in model 1 has suggested that the inhibitory potency of a compound is negatively associated with long length and highly branches of hydrocarbon chains. The impact of the negative association of this index would decrease when the index value has decreased due to the presence of heteroatom. The descriptor [CHI_V_1] is the first order valence molecular connectivity index which the index value decreases with the increase of branches of hydrocarbon chains. This descriptor also considers the presence of heteroatom [58], [59]. Descriptor [CHI_V_1] is positively associated with the inhibitory potency according to the generated 2D-QSAR model 1 and 2, which showed the structures with low branches of hydrocarbon chains and the presence of heteroatom would have good inhibitory activity. Descriptor [CHI_3_P] is the third-order molecular connectivity of the path, which encodes information on the size of 3 adjacent hydrophobic bond fragments with non-hydrogen atoms [59]. The presence of 3 adjacent hydrophobic bond fragments with non-hydrogen atoms is negatively associated with the potency to inhibit β-tryptase. Overall, according to the molecular connectivity index in the generated models, the presence of 1 membered fragment with heteroatom will increase the inhibitory potency of a compound towards β-tryptase. In contrast, a highly branched structure and the presence of 3 adjacent hydrophobic bond fragments containing non-hydrogen atoms will reduce the inhibitory potency of a compound.

Descriptor [E_EDJ_equ] indicates the molar volume and molar refractivity of a compound. This descriptor can discriminate isomers and determine the bulkiness of a model and the position of a heteroatom. The index value increases when heteroatom is positioned at the centre of the structure [60], [61]. The negative association of descriptor [E_EDJ_equ] suggested that high molar volume and molar refractivity of a compound structure would reduce the inhibitory potency of a compound, and the further the position of heteroatom from the centre of the compound structure, the higher the inhibitory potency of the compound. Positive association of descriptor [Num_Rings] indicated a high number of base rings contribute to stronger inhibition of a compound towards β-tryptase [62]. The index value of descriptor [CIC] increases with a higher number of carbon atoms but decreases when the number of hydrogen atoms becomes smaller [63]. The negative contribution of this descriptor in the generated models indicated compound with less carbon in the structure, or a smaller size molecule is preferable, and an unsaturated bond is preferable over a saturated bond.

Analysis from molecular docking for the native ligand of 4A6L, compound 11e, and compound 1a matched with the results from the three 2D-QSAR models. Descriptor ECFP_6:-17546122 showed the presence of a carbonyl group attached to a ring structure is favourable for inhibition activity, and this structure has been found in the native ligand and compound 11e, which the structure has formed a conventional hydrogen bond to the Gly 237 residue of the protein. The NH2 of GLY237 has been reported to form a hydrogen bond with carbonyl oxygen [48], [49]. In contrast, the same structure is not found in compound 1a. Besides, according to the results from 2D-QSAR models, inhibition activity is favourable at low branches of structure and the presence of heteroatom located further from the centre of the structure. The amine groups in the native ligand, compound 11e, and compound 1a are present in the low branched hydrocarbon chain. However, only the amine group of native ligand and compound 11e is located further from the centre of the structure, and their amine group has formed a conventional hydrogen bond to ASP207 and SER208. According to past studies, ASP207 and SER208 are able to form a hydrogen bond with the amidino or amine group [21], [47].

### 3.3 Molecular dynamics (MD) simulation

In MD simulation, results from molecular docking were validated, and the stability of the docked complex was determined. Compound 11e was the most active β-tryptase inhibitor reported, and it consists of structures that are favourable for β-tryptase inhibition according to the 2D QSAR study. In addition, compound 11e also showed good binding affinity and interaction profile in molecular docking analysis. Thus, the docked 4A6L-compound 11e was selected for MD simulation.

Root Mean Square Deviation (RMSD) of the docked complex was plotted as a function of time (Figure 3). The RMSD of 4A6L chain A backbone is at an average of 0.312 and SD of 0.011, and the value is below 2Å. This showed that the simulation system has equilibrated and no large conformational changes in protein throughout the simulation period [42], [64]. RMSD of compound **11e** aligned on protein 4A6L chain A is at an average of 0.437, with an SD of 0.058. These values determine the stability of the compound with respect to the 4A6L chain A. The compound 11e with the RMSD value is only slightly higher than the RMSD of 4A6L chain A, and this shows that the ligand is likely intact with its initial binding site[64].

**Figure 3:**
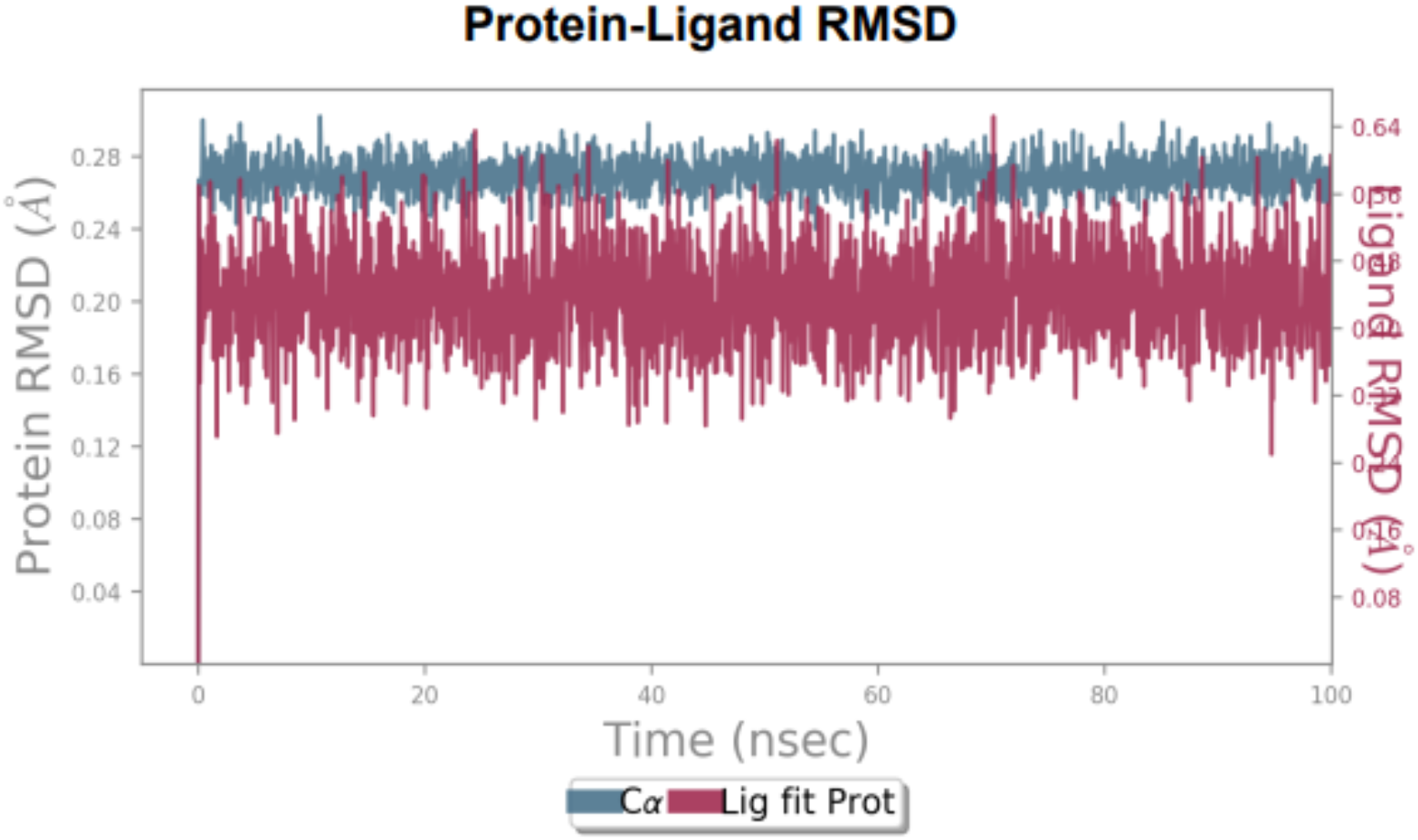
RMSD of 4A6L and compound 11e during MD simulation

Root Mean Square Fluctuation (RMSF) indicates the fluctuations of each residue during the simulation [43]. Protein and ligand in a complex with a lower RMSF value are considered less flexible and are stable complexes [65]. A low RMSF value of protein interacting amino acids indicates the binding of the ligand with protein [45]. RMSF value of c-α atoms in between protein 4A6L chain A residues are around 0.16Å to 0.32Å, as shown in Figure 4. The fluctuation is higher at the N- and C-terminal. Protein residues such as His59, Gln105, Ala185, Ile193, Asp207, Ser208, Gln210, Ser213, Ser232, Trp233, Gly234, Glu235, Gly237, Arg243, Pro244, and Tyr247 have been found to contact with the compound 11e. Among these protein residues, Asp207, Ser208, Gln210, Ser213, Ser232, Trp233, and Gly237 are reported as the important residues at the active site from past studies [21], [47]–[49], [52], [53], [66].

**Figure 4:**
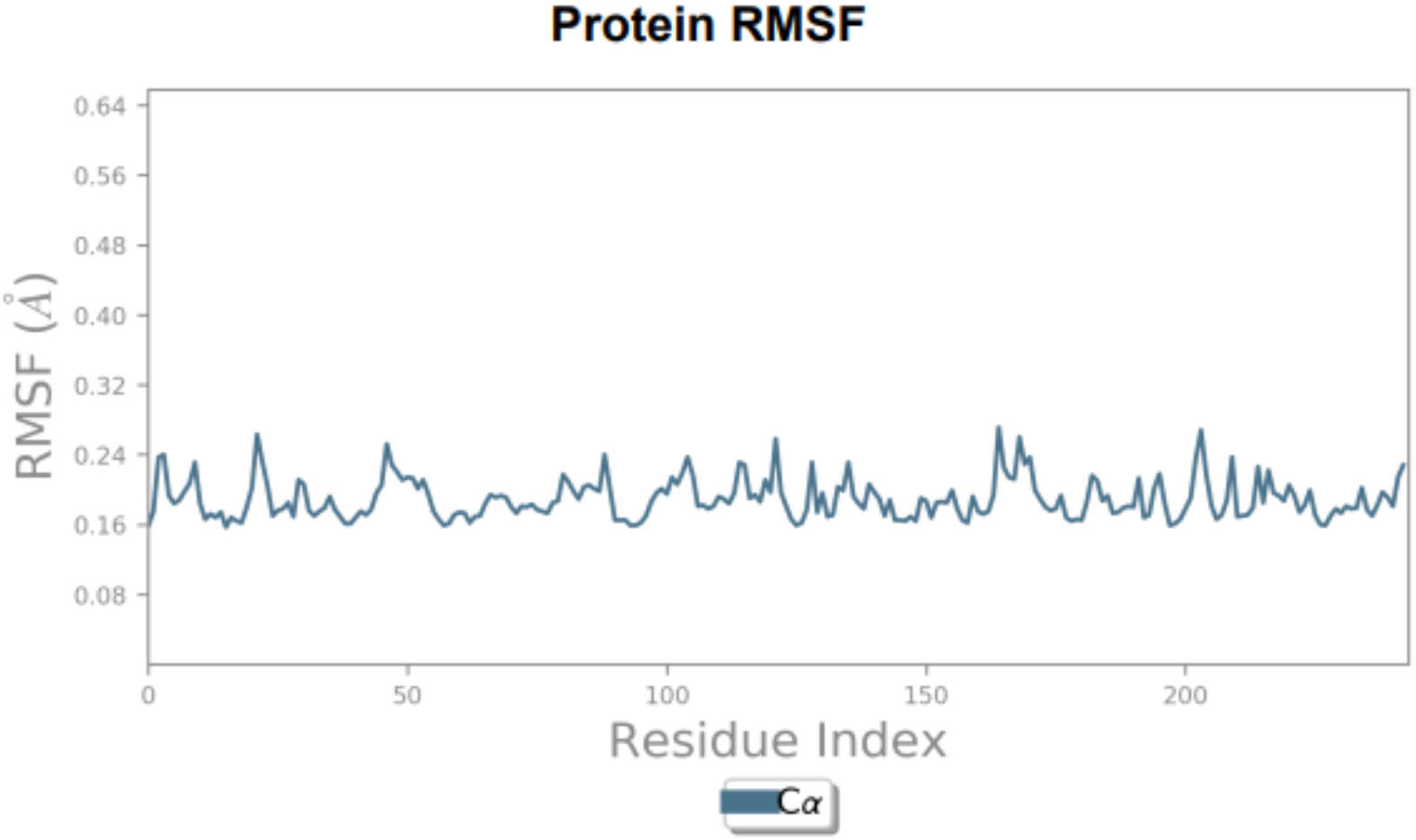
RMSF value of c-α atoms between residues in 4A6L Chain A during MD simulations

RMSF value determines the stability of a compound in the active site (Figure 5). The maximum fluctuation of a few atoms in compound 11e reached 0.4Å and above. This occurred to the terminal atoms of the compound such as atom 7th and atom 43^rd^, these atoms are not involved in any interaction [67]. On the other hand, most of the atoms with RMSF at around or below 3Å. This showed that a large part of the compound 11e has lower fluctuation and the compound is attached to the active site of the protein [45]. Figure 6 showed the involvement of atoms 21^st^, 23^rd^, 24^th^, 26^th^, 28^th^, and 34^th^ with the 4A6L catalytic residues at the active site.

**Figure 5:**
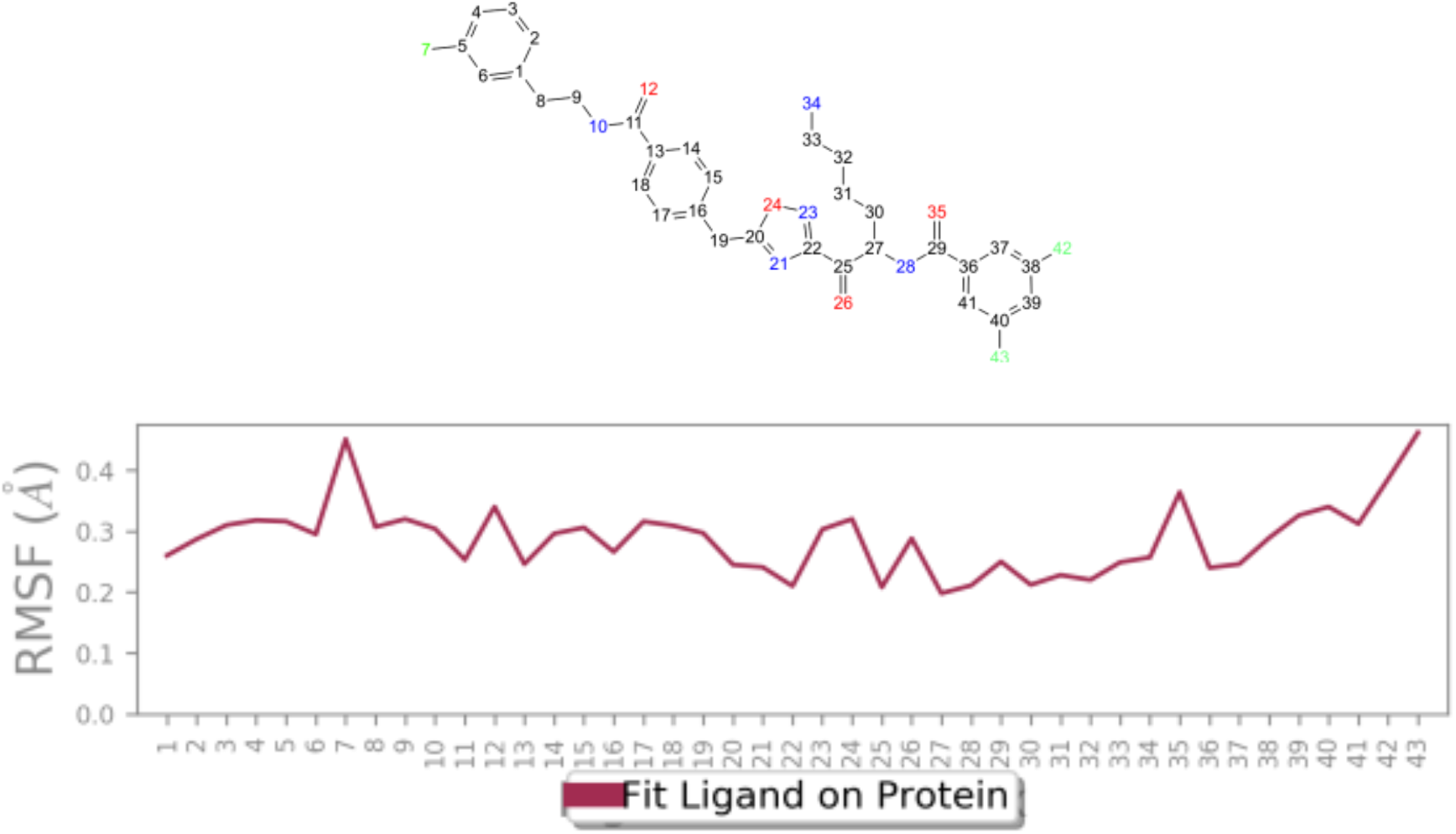
RMSF value of ligand during MD simulations

**Figure 6:**
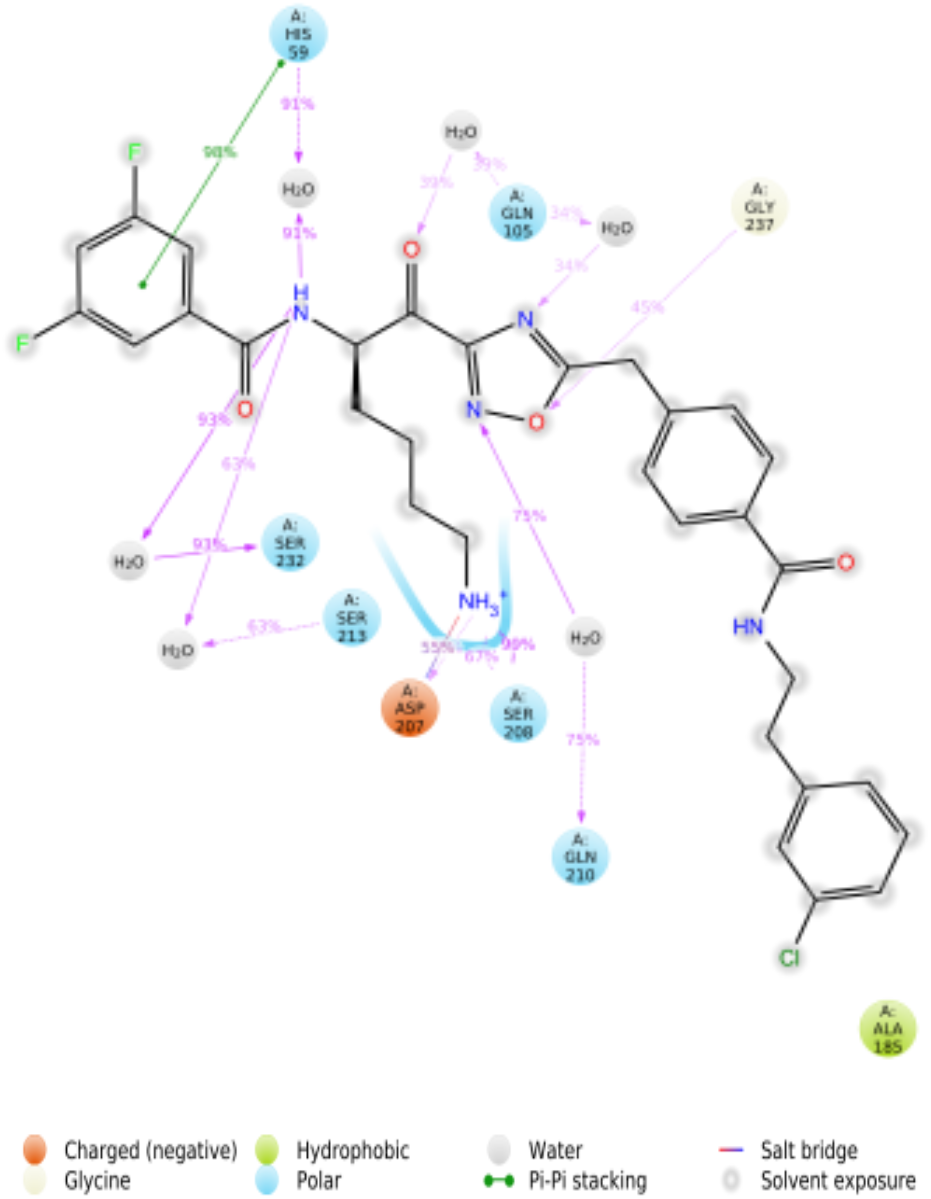
Interaction of compound **11e** with protein 4A6L Chain A from MD simulations

The interaction types of the docked complex identified from the MD simulation included hydrogen bond, hydrophobic interaction, ionic interaction, and water bridges (Figure 7). Hydrogen bonds formed in the interaction are contributed by residues Asp207, Ser208, Gly237, and Gly234. Some residue formed water bridge interactions with compound 11e, which included Ser232, Gln210, Ser213, Gln105, His59, Glu235, Gly237, and Asp207. Residue Asp207, which is one of the catalytic residues at the S1 pocket of the active site of β-tryptase, contributed to the formation of ionic interaction. It has been reported that Asp207 is able to deprotonate its carboxylate side chain to form ionic interaction with basic structure [47]. Hydrophobic interaction is contributed by residue His59, ala185, Ile193, trp233, and tyr247. Table 3 summarised the type and contribution of interactions between key residues of β-tryptase and compound **11e** throughout the MD simulation.

**Figure 7:**
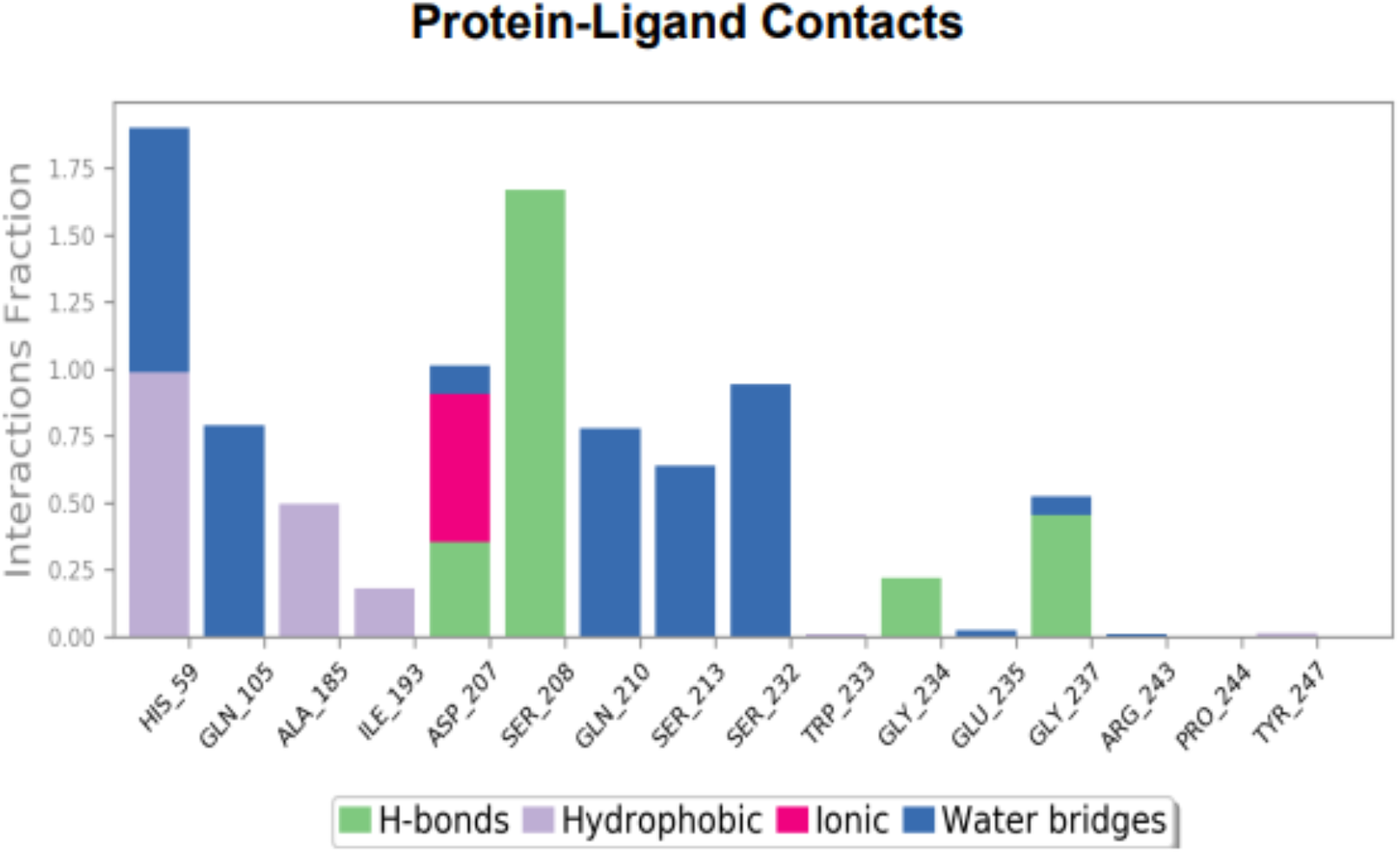
The interaction types of the docked protein 4A6L Chain A-compound 11e complex identified from MD simulation

**Table 3:**
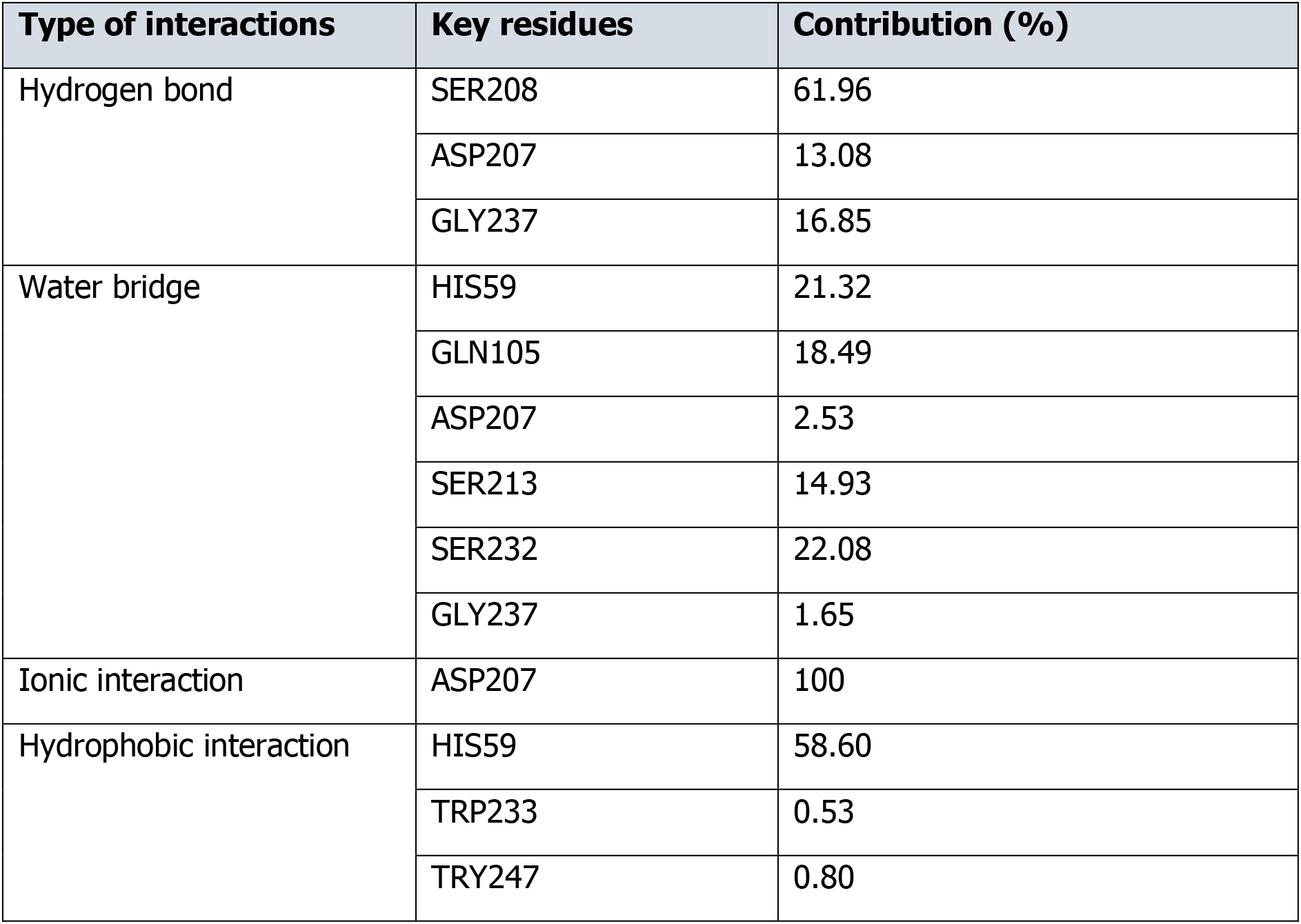
Type and contribution of interactions between key residues of β-tryptase and compound 11e throughout the MD simulation

### 3.4 Pharmacophore modelling

The pharmacophore model generated from 2.5 is shown in Figure 8. There are four pharmacophore feature types in the generated model: aromatic ring feature, hydrogen-bond acceptor feature, ionic interaction feature, and hydrophobic interaction feature. The seven exclusion volumes are kept at the centre of the binding pocket. The pharmacophore model generated and edited is matched with the interaction reported from the MD simulation of compound 11e with 4A6L (Figure 6). At the aromatic ring feature, the structure formed pi-pi stacking interaction with His59 residue. While at the ionic interaction feature, the amino group formed a salt bridge with charged Asp207 residue. Besides, Gly237 residue donated hydrogen to oxygen in the furan ring to form a hydrogen bond, where the hydrogen bond acceptor feature is located. Lastly, hydrophobic interaction with ALA185 residue has occurred near the phenyl chloride structure, where the hydrophobic feature is located.

**Figure 8:**
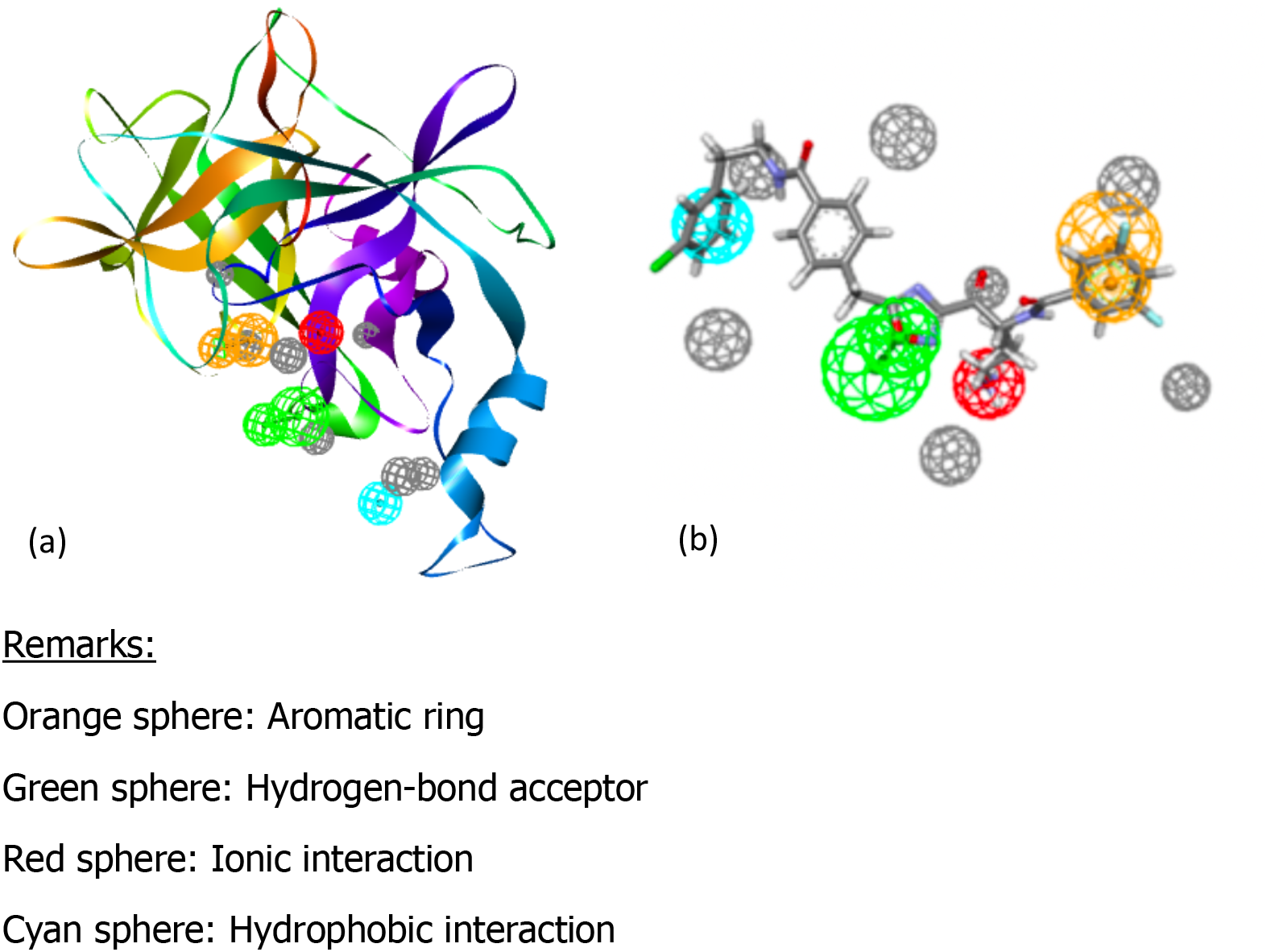
Diagram illustrates the generated structure-based pharmacophore model with compound 11e. (a) Pharmacophore model in 4A6L chain A structure. (b) Pharmacophore model and compound 11e.

The results of the pharmacophore model validation are shown in Table 4. The pharmacophore model had a low hit rate but a moderate yield of actives and moderate sensitivity, with a GH score of 0.46, implying that at least 46% of the returned compounds were not hit randomly from the screening [45]. On the other hand, it has a good EF of 57.15, showing that the model has 57.15 times the possibility of detecting active compounds over inactive compounds [45]. Besides, the pharmacophore model has very high accuracy and specificity, which are 99.03% and 99.90%, respectively. The ROC curve has been generated to predict the performance of the pharmacophore model (Figure 9). The AUC is 0.722, indicating that the pharmacophore model has a fair performance [46], [68].

**Table 4:** Validation result of Pharmacophore model

|  |  |  |
| --- | --- | --- |
|  | Total compounds in the dataset (D) | 8701 |
|  | Total number of actives in the dataset (A) | 87 (= 1% of the total dataset) |
| 1 | Total hits | 21 |
| 2 | Total true positive | 12 |
|  | Total false negative | 66 |
|  | Total false positive | 9 |
|  | Total true negative | 8605 |
| 3 | Enrichment factor (EF) | 57.15 |
| 4 | Güner-Henry Score (GH score) | 0.46 |
| 5 | Yield of actives (%) | 57.14 |
| 6 | Accuracy (%) | 99.03 |
| 7 | Sensitivity (%) | 13.79 |
| 8 | Specificity | 99.90 |

**Figure 9:**
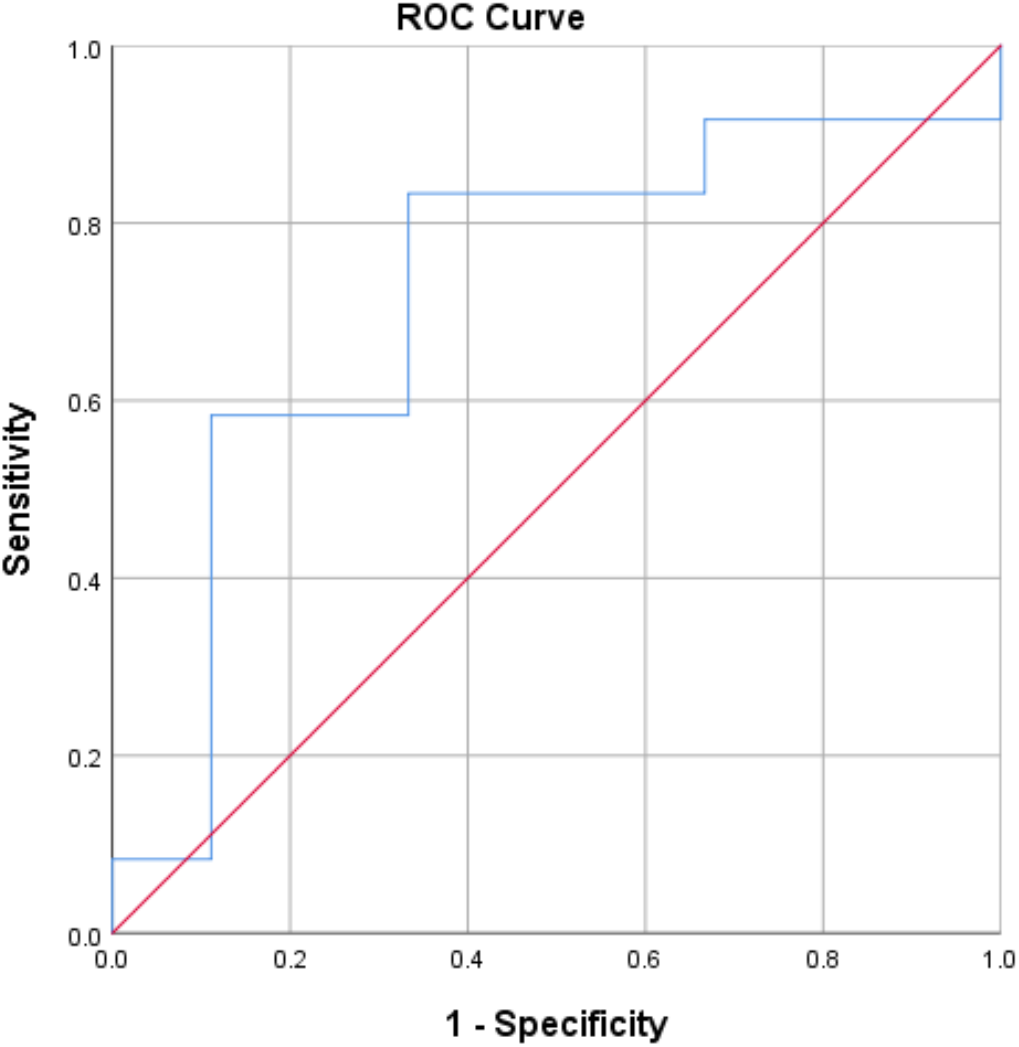
ROC Curve of the generated pharmacophore model with AUC of 0.722.

## 4. Conclusion

2D QSAR modelling of α-keto-[1,2,4]-oxadiazoles scaffold-based derivatives was used to create a model for the β-tryptase inhibitor. The GFA technique created a QSAR equation with good quality and predictive capability. Compounds with structure of molecular descriptors [ECFP_6: 76543811], [ECFP_6: −522073680], and [ECFP_6: 1205550831] are predicted to enhance the inhibition of β-tryptase. Besides, the inhibitory activity of compounds is predicted to increase with the presence of membered fragments with heteroatom located further from the centre of the compounds’ structure. Furthermore, compounds with a high number of base rings, fewer carbon atoms in the structure, and smaller molecular size, as well as the presence of unsaturated bonds are also able to improve their inhibition activity towards β-tryptase. On the other hand, molecular docking study revealed that the most active compound, 11e has lower binding energy than the co-crystalised β-tryptase native inhibitor. The binding pattern of compound 11e is similar to the native ligand. MD simulation confirmed the stable binding of compound 11e with protein 4A6L Chain A. The results are consistent with the findings of [32], in which compound 11e was reported with the highest inhibitory potency. In this study, the pharmacophore model generated with features matched the interactions of compound 11e with protein 4A6L Chain A obtained from MD simulation. Hence, the information offered by this study could be used to develop a more effective β-tryptase inhibitor for the treatment of vascular leakage during DENV infection.

## Acknowledgments

The authors would like to thank the members of Cell Signalling Laboratory, Faculty of Medicine and Health Sciences, Universiti Putra Malaysia, for providing supports for this study. The authors would also like to thank International Islamic University Malaysia as well as Universiti Kebangsaan Malaysia (UKM) for providing Discovery Studio^®^ 3.1 software. The authors are also thankful to Atta-ur-Rahman Institute for Natural Product Discovery (AuRIns) and the Faculty of Applied Science, Universiti Teknologi Mara for support in molecular dynamics simulation.

## List of Abbreviations

DENV: Dengue virus
DF: Dengue Fever
DHF: Dengue hemorrhagic fever
DSS: Dengue shock syndrome
MC: Mast cell
CADD: Computer-aided drug design
QSAR: Quantitative structure-activity relationship
MD: Molecular dynamics
PM: Pharmacophore modelling
DS: Discovery Studio^®^ 3.1
PDB: Protein Data Bank
RMSD: Root mean square deviation
Ki: Inhibition constant
GFA: Genetic Function Approximation
ROC: Receiver operating characteristics
AUC: Area under the curve
EF: Enrichment factor
GH: Güner-Henry
RMSF: Root mean square fluctuation

**\*This pre-print is a manuscript submitted to the Journal of Molecular Graphics and Modelling**

## Notes

### Competing Interest Statement

The authors have declared no competing interest.

