## Supplemental Table 1 for "Insight parameter drug design for human β-tryptase inhibition integrated molecular docking, QSAR, molecular dynamics simulation, and pharmacophore modelling studies of α-keto-[1,2,4]-oxadiazoles"

**Table S1:** The structures of the data set and its biological activities from [19]

| Compounds | Structure | Ki (nM) | pKi (M) |
|---|---|---|---|
| 1a | 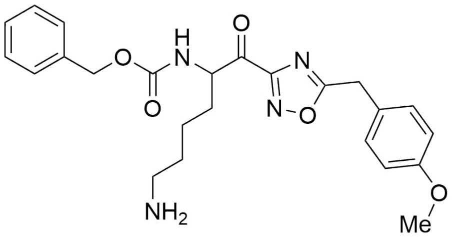 | 770 | 6.11 |
| 1b | 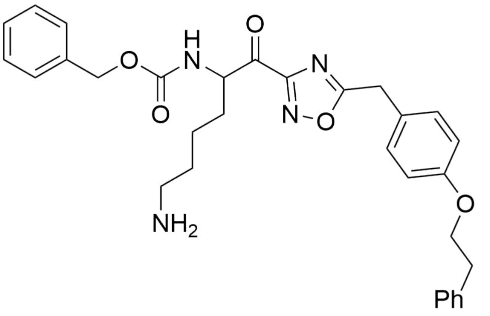 | 12 | 7.92 |
| 6a | 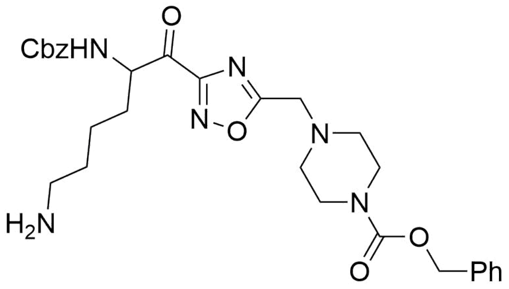 | 470 | 6.33 |
| 6b | 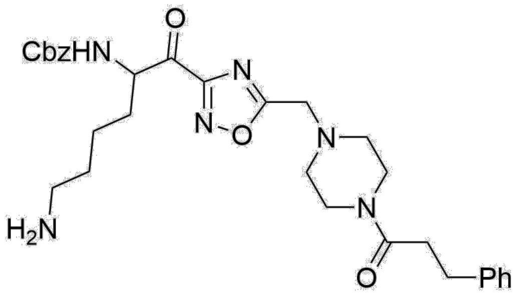 | 230 | 6.64 |
| 6c | 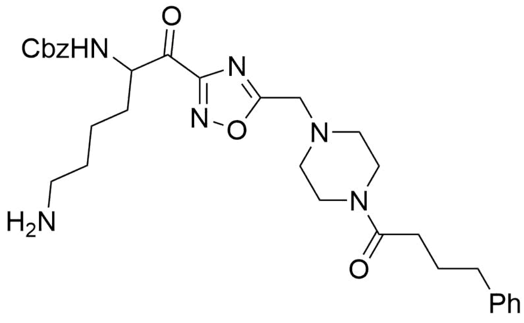 | 34 | 7.47 |

| | | | |
|---|---|---|---|
| 6d | 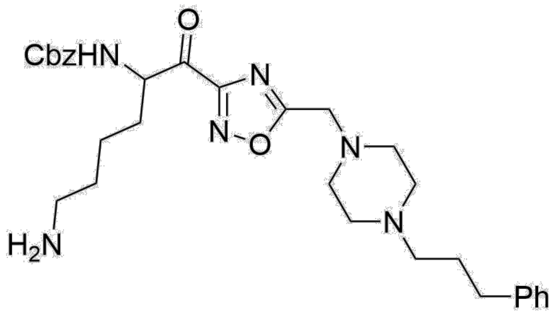 | 11 | 7.96 |
| 6e | 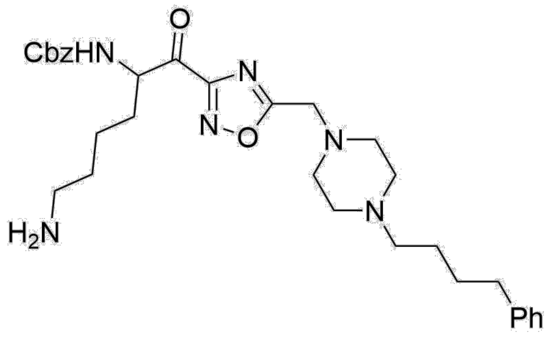 | 5.8 | 8.24 |
| 10a | 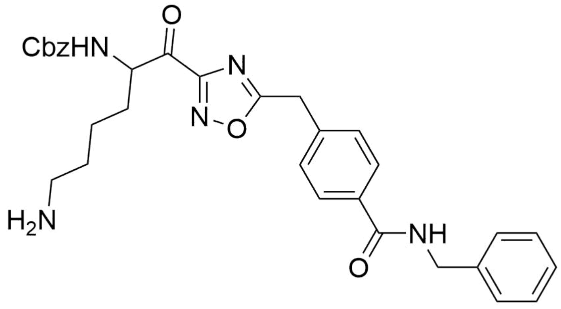 | 37 | 7.43 |
| 10b | 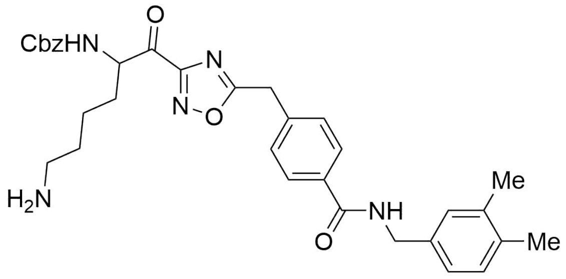 | 39 | 7.41 |
| 10c | 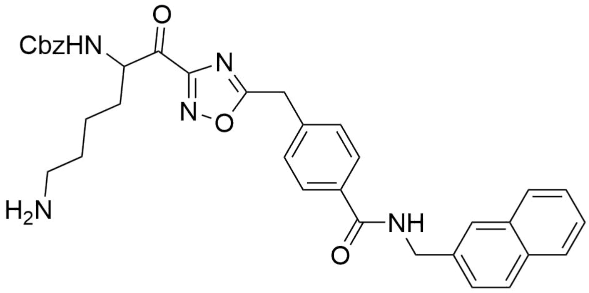 | 11 | 7.96 |

| | | | |
|---|---|---|---|
| 10d | 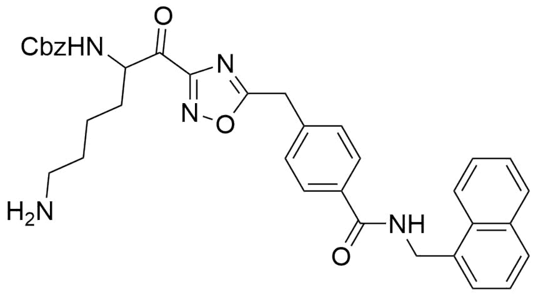 | 13 | 7.89 |
| 10e | 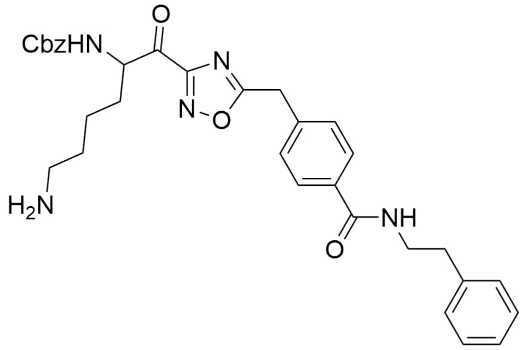 | 3.2 | 8.49 |
| 10f | 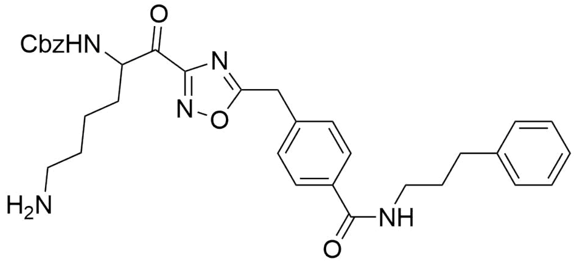 | 3.0 | 8.52 |
| 10g | 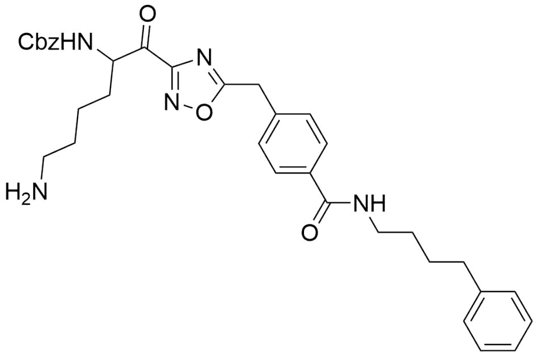 | 60 | 7.22 |
| 10h | 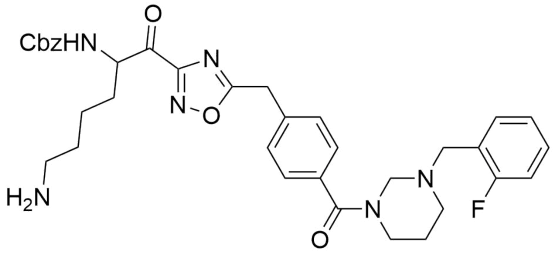 | 140 | 6.85 |

| | | | |
|---|---|---|---|
| 10i | 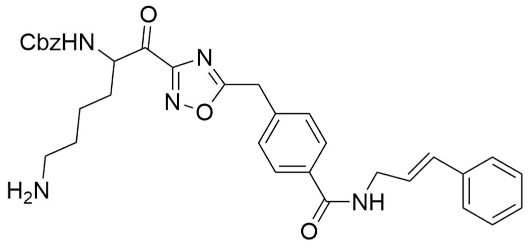 | 3.6 | 8.44 |
| 10j | 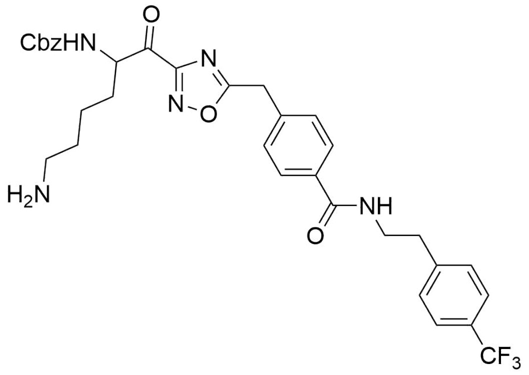 | 220 | 6.66 |
| 10k | 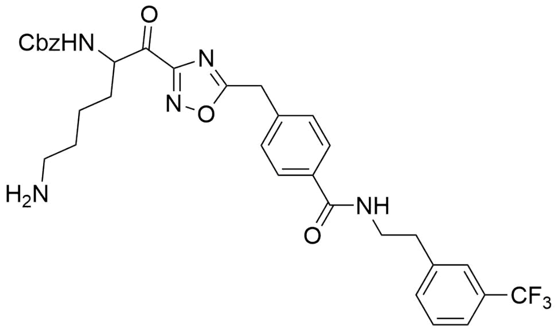 | 8.2 | 8.09 |
| 10l | 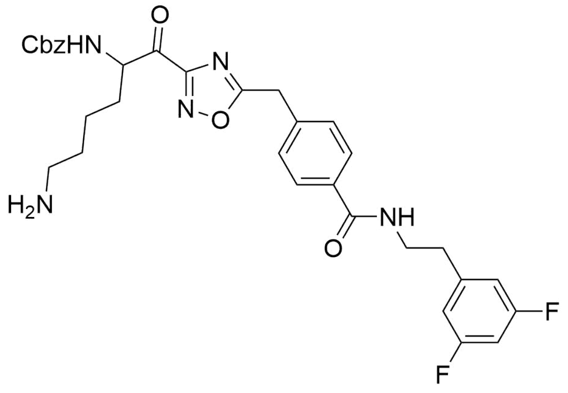 | 3.5 | 8.46 |

| | | | |
|---|---|---|---|
| 10m | 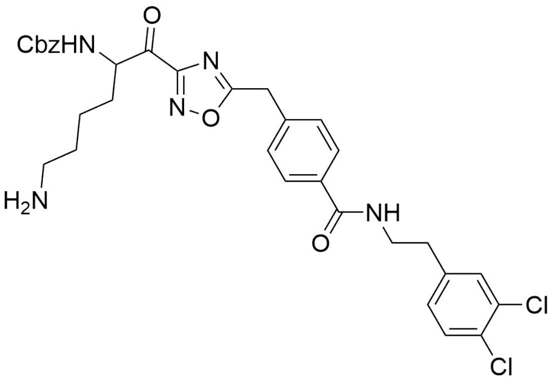 | 16 | 8.00 |
| 10n | 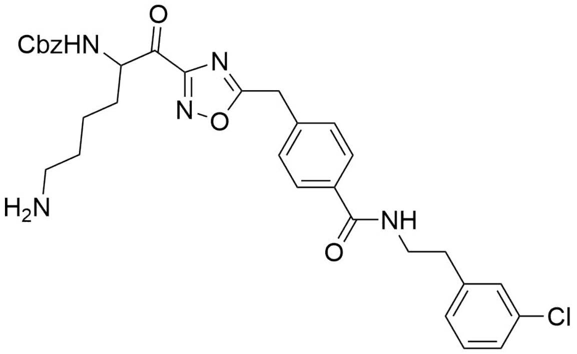 | 1.8 | 8.74 |
| 10o | 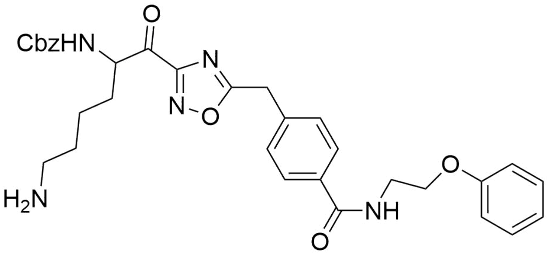 | 2.9 | 8.54 |
| 10p | 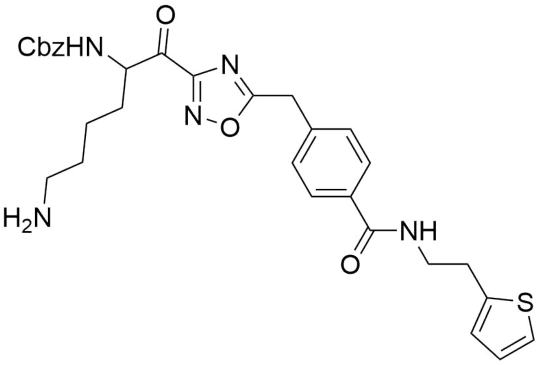 | 2.4 | 8.62 |
| 10q | 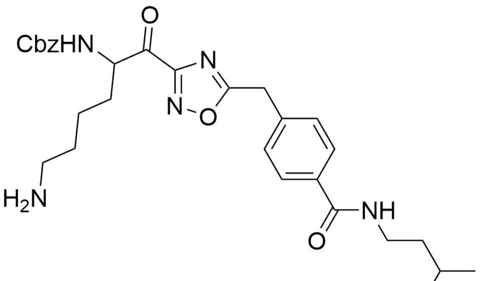 | 9.9 | 8.00 |

| | | | |
|---|---|---|---|
| 10r | 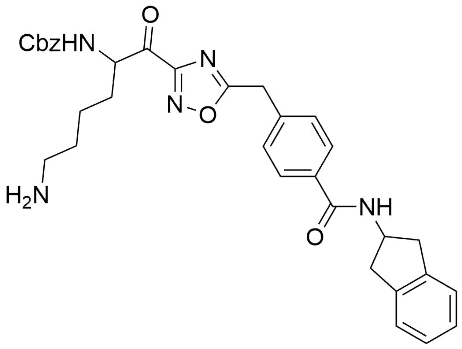 | 3.7 | 8.43 |
| 11a | 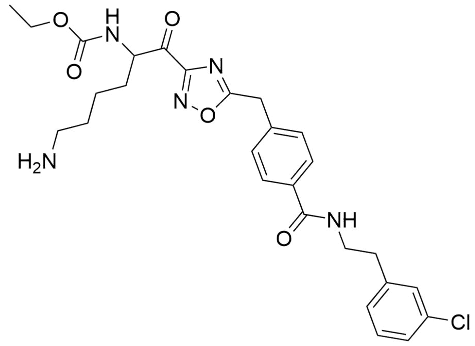 | 1.9 | 8.72 |
| 11b | 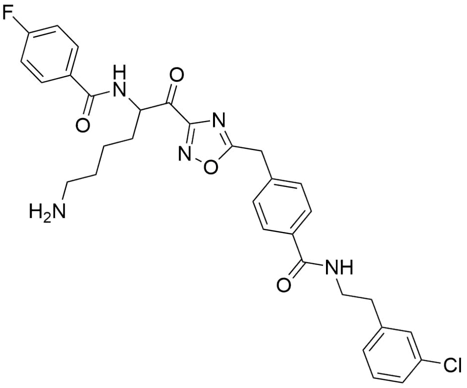 | 1.8 | 8.74 |
| 11c | 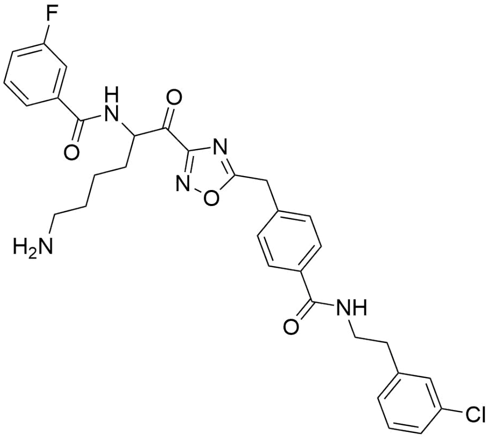 | 1.7 | 8.77 |

| | | | |
|---|---|---|---|
| 11d | 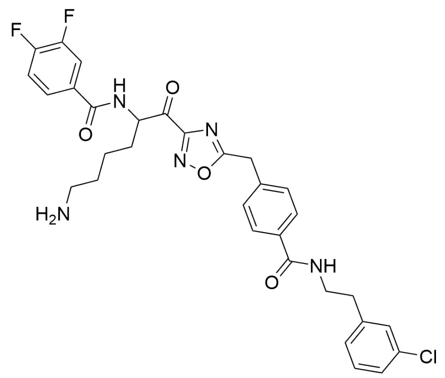 | 1.7 | 8.77 |
| 11e | 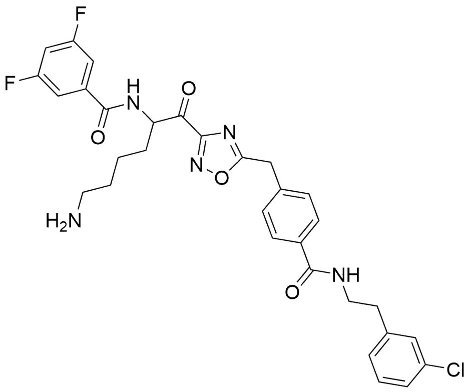 | 1.5 | 8.82 |
| 11f | 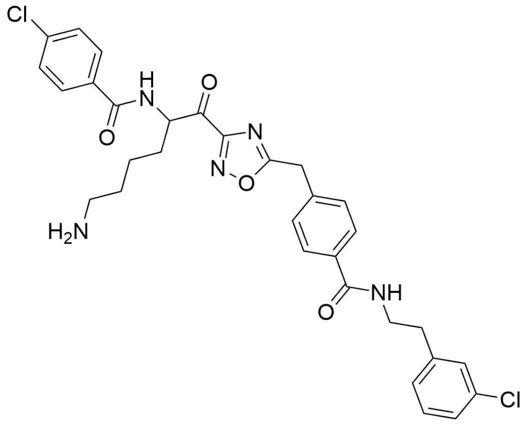 | 1.9 | 8.72 |
| 11g | 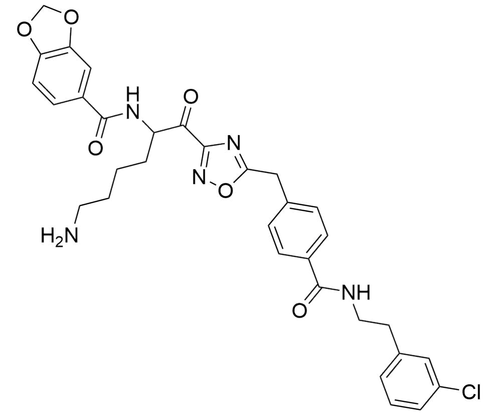 | 2.1 | 8.68 |

| | | | |
|---|---|---|---|
| 11h | 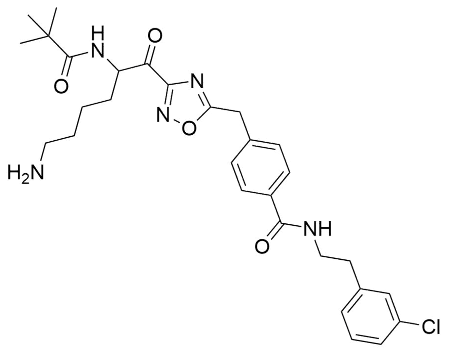 | 2.0 | 8.7 |
| 12a | 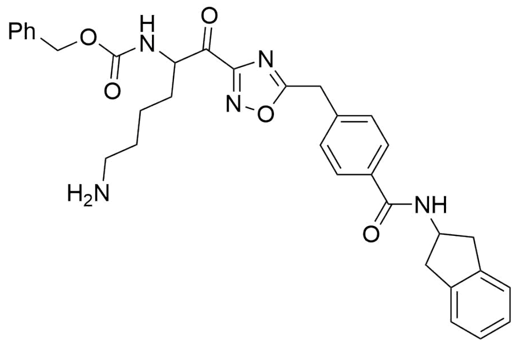 | 3.7 | 8.43 |
| 12b | 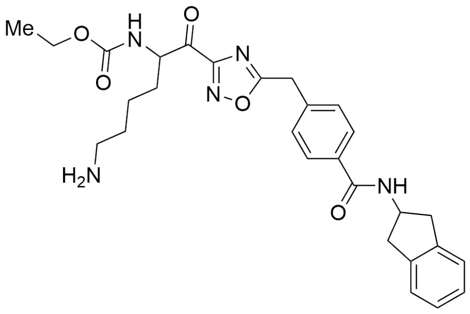 | 2.8 | 8.55 |
