## Supplemental Table 4 for "Insight parameter drug design for human β-tryptase inhibition integrated molecular docking, QSAR, molecular dynamics simulation, and pharmacophore modelling studies of α-keto-[1,2,4]-oxadiazoles"

**Table S4:** Descriptors [ECFP_6: 765434811], [ECFP_6:-522073680] and [ECFP_6:-175146122] found in the structure of β-tryptase inhibitors from past studies.

| Source | Inhibitors | Inhibitory activities |
|---|---|---|
| [40] | Nafamostat 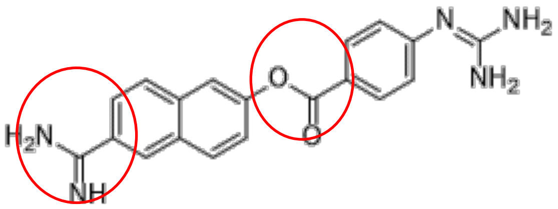 | Ki 95.3pM |
| [40] | Gabexate 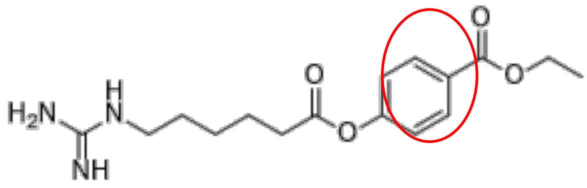 | Ki 95.1nM |
| [36] (co-crystallised ligand of PDB ID 4A6L) | 1-{3-[1-({5-[(2-fluorophenyl)ethynyl]furan-2-yl} carbonyl)Piperidin-4-yl]phenyl}methanamine 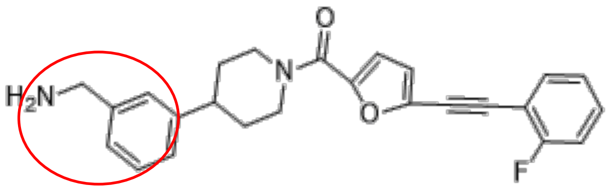 | Ki = 10nM |
| [41] (co-crystallised ligand of PDB ID 5F03) | (5~{S})-5-[[3-(aminomethyl)phenoxy]methyl]-3-[3-[2-(2-chloranylpyridin-3-yl)ethynyl]phenyl]-1,3-oxazolidin-2-one 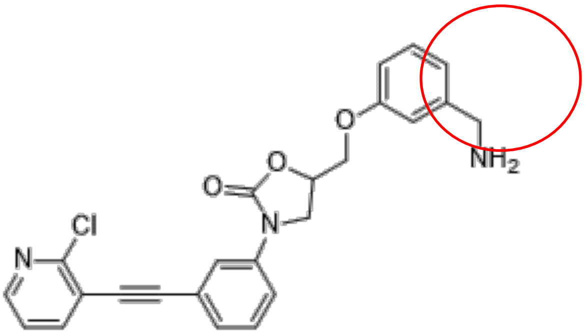 | Ki = 50nM |

| | | |
|---|---|---|
| [34]<br>(co-crystallised ligand of PDB ID 2BM2) | 1-[3-(1-{[5-(2-Phenylethyl)Pyridin-3-yl]Carbonyl}Piperidin-4-yl)Phenyl]Methanamine<br>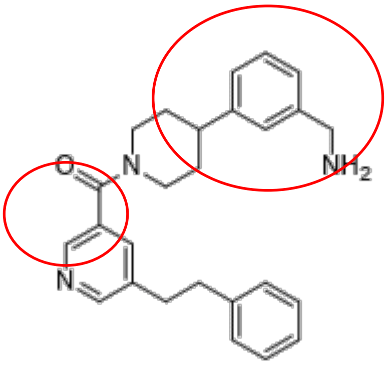 | Ki = 15nM |
| [42]<br>(co-crystallised ligand of PDB ID 2FS9) | Ethyl{(1S)-5-amino-1-[(5-{4-[(2,3-Dihydro-1H-Inden-2-ylamino)Carbonyl]Benzyl}-1,2,4-Oxadiazol-3-yl)Carbonyl]Pentyl}Carbamate<br>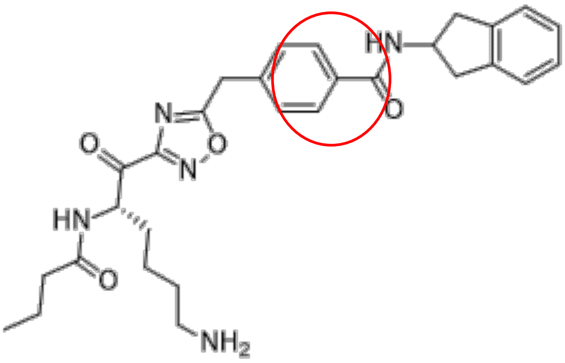 | Ki 2.8nM |
| [43] | APC-366<br>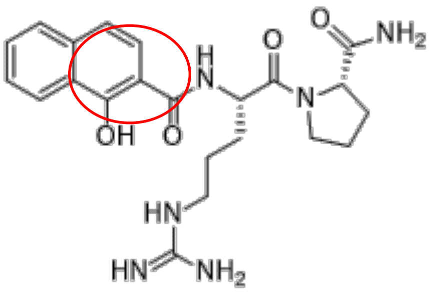 | IC50 300nM |
| [44] | Active site bridging inhibitors of human tryptase | Ki 15nM |

| | | |
|---|---|---|
| | 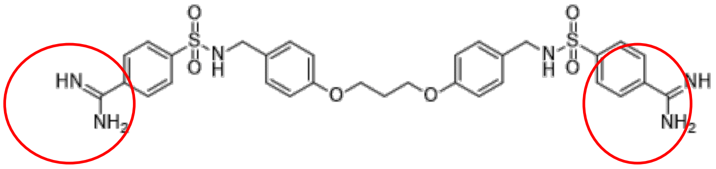 | |
| [45] | Compound 4e 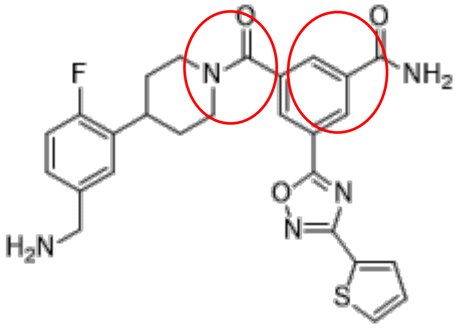 | Ki = 1.3nM |
| [46] | Compound 1.14 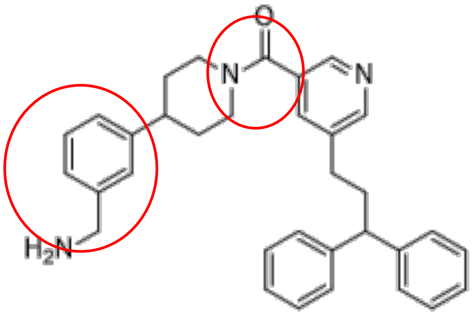 | Ki 1.5nM |
| [47] | BABIM 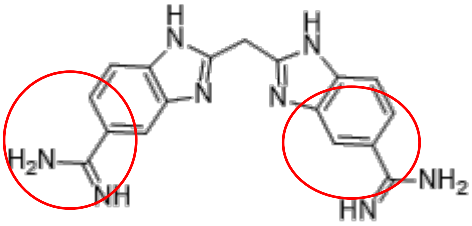 | Ki = 1.8nM |
