## Supplemental table 3 for "Insight parameter drug design for human β-tryptase inhibition integrated molecular docking, QSAR, molecular dynamics simulation, and pharmacophore modelling studies of α-keto-[1,2,4]-oxadiazoles"

**Table S3:** Experimental and predicted inhibition activities (pKi, M) of the compounds from three selected 2D-QSAR models

| | Compounds | Experimental pKi (M) | Predicted pKi (M) | | |
|---|---|---|---|---|---|
| | | | Model 1 | Model 2 | Model 3 |
| Training set | 1b | 7.92 | 7.99 | 7.95 | 7.95 |
| | 6a | 6.33 | 6.40 | 6.30 | 6.21 |
| | 6b | 6.64 | 6.99 | 6.93 | 7.12 |
| | 6c | 7.47 | 7.99 | 7.95 | 7.95 |
| | 6e | 8.24 | 7.82 | 7.98 | 7.88 |
| | 10a | 7.43 | 7.71 | 7.68 | 7.45 |
| | 10b | 7.41 | 7.33 | 7.31 | 7.30 |
| | 10c | 7.96 | 7.97 | 7.98 | 8.01 |
| | 10d | 7.89 | 7.99 | 7.92 | 8.06 |
| | 10e | 8.49 | 8.52 | 8.52 | 8.56 |
| | 10f | 8.52 | 8.52 | 8.52 | 8.56 |
| | 10h | 6.85 | 6.72 | 6.89 | 6.88 |
| | 10i | 8.44 | 8.02 | 7.95 | 7.97 |
| | 10j | 6.66 | 6.72 | 6.89 | 6.88 |
| | 10k | 8.09 | 7.96 | 7.78 | 7.92 |
| | 10l | 8.46 | 7.99 | 7.95 | 7.95 |
| | 10m | 8.00 | 8.01 | 7.97 | 7.97 |
| | 10n | 8.74 | 9.07 | 9.12 | 9.00 |
| | 10o | 8.54 | 8.52 | 8.52 | 8.56 |
| | 10r | 8.43 | 8.46 | 8.48 | 8.43 |
| | 11a | 8.72 | 8.68 | 8.72 | 8.78 |
| | 11b | 8.74 | 8.86 | 8.84 | 8.72 |
| | 11c | 8.77 | 8.68 | 8.72 | 8.78 |
| | 11d | 8.77 | 8.68 | 8.72 | 8.78 |
| | 11f | 8.72 | 8.86 | 8.84 | 8.72 |
| | 11g | 8.68 | 8.47 | 8.48 | 8.47 |
| | 11h | 8.70 | 8.68 | 8.72 | 8.78 |
| Test set | 1a | 6.11 | 6.72 | 6.89 | 6.88 |
| | 6d | 7.96 | 7.82 | 7.98 | 7.88 |

| | | | | | |
|---|---|---|---|---|---|
| | 10g | 7.22 | 7.99 | 7.95 | 7.95 |
| | 10p | 8.62 | 8.86 | 8.84 | 8.72 |
| | 10q | 8.00 | 8.68 | 8.72 | 8.78 |
| | ★ 11e | 8.82 | 8.52 | 8.52 | 8.56 |
| | 12b | 8.55 | 8.09 | 8.08 | 8.22 |
