## Supplemental table 2 for "Insight parameter drug design for human β-tryptase inhibition integrated molecular docking, QSAR, molecular dynamics simulation, and pharmacophore modelling studies of α-keto-[1,2,4]-oxadiazoles"

**Table S2:** Docking result analysis based on binding interaction of the compounds with 4A6L residues and their CDOCKER energy.

| Compounds | -CDOCKER Energy | Interacting residues | | | | | | |
|---|---|---|---|---|---|---|---|---|
| | | ASP207 | Ser208 | Ser213 | Ser232 | Gly237 | His59 | Gln105 |
| β-tryptase inhibitor | 23.9672 | \ | \ | | | \ | | |
| 11e_1.5nM | 40.9846 | \ | \ | | | \ | | \ |
| 1a_770nM | 36.3865 | | \ | | | \ | \ | |
| 1b_12nM | 46.6222 | \ | \ | | | \ | \ | \ |
| 6c_34nM | 44.3164 | \ | \ | | | \ | \ | \ |
| 6d_11nM | 37.7296 | | | \ | | \ | | |
| 6e_5.8nM | 40.5026 | \ | \ | | | \ | | |
| 10a_37nM | 51.917 | | \ | \ | \ | \ | | |
| 10b_39nM | 48.8915 | \ | \ | \ | \ | \ | \ | |
| 10c_11nM | 35.6865 | | \ | \ | \ | | | \ |
| 10d_13nM | 35.9447 | \ | \ | \ | | \ | \ | |
| 10e_3.2nM | 49.0806 | | | | \ | \ | \ | \ |
| 10f_3.0nM | 35.0137 | | \ | \ | | | | \ |
| 10i_3.6nM | 31.747 | | | \ | | \ | \ | |
| 10k_8.2nM | 50.3211 | \ | \ | | | \ | | \ |
| 10l_3.5nM | 45.2573 | | | | | \ | \ | |
| 10m_10nM | 27.2416 | | | | | \ | | \ |
| 10n_1.8nM | 42.9754 | | \ | \ | \ | \ | | \ |

| | | | | | | | | |
|---|---|---|---|---|---|---|---|---|
| 10o_2.9nM | 42.9523 | | \ | \ | \ | \ | \ | \ |
| 10p_2.4nM | 47.3897 | | \ | | \ | \ | \ | |
| 10q_9.9nM | 42.2149 | | | | | | | \ |
| 10r 3.7nM | 24.7219 | | | \ | | | \ | \ |
| 11a_1.9nM | 46.5288 | | | \ | | \ | \ | |
| 11b_1.8nM | 43.8614 | \ | \ | | | | \ | |
| 11c_1.7nM | 42.8161 | \ | \ | \ | | \ | \ | |
| 11d_1.7nM | 32.6984 | | \ | \ | | \ | | |
| 11f_1.9nM | 40.1707 | | | \ | | | \ | |
| 11g_2.1nM | 21.7144 | \ | \ | | | \ | \ | \ |
| 11h_2.0nM | 50.3727 | \ | \ | | | \ | \ | \ |
| 12b_2.8nM | 27.4054 | \ | \ | \ | \ | \ | | |
| 10g_60nM | 42.6587 | | | \ | | \ | \ | |
| 6a_470nM | 40.0379 | \ | | \ | | \ | \ | |
| 6b_230nM | 35.4884 | \ | \ | \ | | \ | \ | |
| 10h_140nM | 37.7629 | | | \ | | | | \ |
| 10j_220nM | 39.1587 | | | \ | \ | \ | \ | |
